# MEG Oscillatory and Aperiodic neural dynamics contribute to different cognitive aspects of short-term memory decline through lifespan

**DOI:** 10.1101/2021.03.02.433594

**Authors:** Kusum Thuwal, Arpan Banerjee, Dipanjan Roy

## Abstract

Neural communication or signal transmission in the brain propagates via distinct oscillatory frequency bands. With aging, the communication mediated by these frequency bands is hindered by noise, which arises from the increased stochastic variability in the baseline neural spiking. This increase in noise measured as 1/f power-law scaling reflects the global background noise and is often linked to impaired cognition in different tasks. In this study, we quantified the 1/f slope and intercept of MEG brain signal as a putative marker of neural noise and examined its effect on cognitive and metacognitive measures. We hypothesize that as neural communication becomes noisier with age, it impacts global information processing, whereas specific periodic features mediate local aspects of cognition. Using recently proposed parametric Fooof model, we first characterised the normative pattern of periodic and aperiodic features (temporal dynamics) across the lifespan, modelled via spectral peaks (Central frequency, power, bandwidth) and 1/f noise activity (slope and intercept) respectively. Secondly, how this Resting-State (RS) baseline shift in temporal dynamics of the signal is associated with various aspects of visual short-term memory (VSTM). Our results suggest that age-associated global change in noisy baseline affects global information processing and crucially impacts the oscillatory features, which relates to more local processing and selective behavioural measures in the VSTM task. Moreover, we suggest that the task-related differences observed across age groups are due to the baseline shift of periodic and aperiodic features.

**Significant statement:** Aging is accompanied by the decline in cognitive functions and age itself is a major risk factor for Alzheimer’s Disease and other neurological conditions. Our study provides MEG 1/f aperiodic and periodic markers across the healthy adult lifespan and shows that different frequency bands and their spectral features mediate age-related changes across different brain regions, in multiple cognitive and metacognitive domains, which not only provides us with a better understanding of the aging process but would also help in better prevention of cognitive impairments. A clear characterization of the association between baseline MEG temporal dynamics, healthy aging and cognition, is established in this study.

## INTRODUCTION

It has recently become more evident that spontaneous oscillations in the electrical potential are not just brain activity engulfed with noise, rather the local as well global change in spontaneous dynamics may index key aspects of behavioural response associated with healthy aging process (Bishop et al., 2010; Foster et al., 2015; Sahoo et al., 2020). The brain oscillations emerging in the spontaneous state based on the underlying intrinsic neuronal coupling even in the absence of goal-directed tasks allow us to quantify normative patterns of brain dynamics and it’s marked alterations through the lifespan to the extent to even capture salient aspects of cognitive functions in the domain of attention, perception, and memory processing (Buzsáki, G. 2006). In principle, neural oscillation ideally should represent the rhythmic activity of the signal arising from different sensors or underlying brain sources, however, non-oscillatory/aperiodic component (1/f noise) almost always pervasively co-exists with the oscillatory components. This background noise has been modelled in the literature as a characteristic 1/f component (slope and offset of the ongoing oscillatory power) and found to be very dynamic in nature (Grigolini et al., 2009; Haller et al., 2018; Voytek et al., 2015). One possible mechanism underlying this dynamical change is an alteration in the underlying neuronal population spiking stats and the relationship of the spiking dynamics on the underlying change in the excitation-inhibition balance which is primarily reflected as an increased baseline activity. A large majority of the works till this day has used predefined canonical frequency bands to examine rhythmic activities without giving substantial amounts of considerations to the aperiodic signal which strongly impacts the oscillatory signal measured from various sensors and brain regions. Therefore, investigation of neural oscillations using the traditional signal processing analysis might not represent the true oscillation rather the dynamically changing slope or offset within the 1/f signal as was suggested recently (Donoghue et al., 2020; Ouyang et al., 2020). Physiological aging has been characterized as a progressive change in Oscillatory power, central frequency, and functional connectivity between relevant brain areas indexing decline in coordination dynamics during resting-state and task conditions. While there is general consensus among findings what typically constitute neural correlates of age-associated oscillatory changes e.g., (i) slowing down of central frequency of alpha band(7–12 Hz), and (ii) global increase of beta (13–30 Hz) and theta band power (4–8 Hz) (Klimesch et al., 1999), there are also noticeable discrepancies among the existing studies. For instance, studies have reported mixed evidence for the difference in the theta band power associated with healthy aging. While many of the earlier studies have reported that theta band power tends to increase (Cummins et al., 2007; Klass et al., 1995) some of the later studies tend to suggest the opposite (Stomrud et al., 2010) with age. For the alpha band, while few studies have reported no change (Aurlien et al., 2014; Scally et al., 2018), others, have reported a decrease in power (Ishii et al.,2017).

Hence, it is often notoriously difficult to reconcile those age-associated oscillatory findings during spontaneous activity and trusting power changes in the relevant frequency band were estimated accurately. . One possible reason for this inconsistency might be the mixing of oscillatory power with the background 1/f activity, which was not taken into sufficient consideration by most of the studies.

The aperiodic 1/f noise of the background activity also attempts to account for changes observed associated with aging, for example, to ascertain memory (Nyberg et al, 2012), shift of sustained attention(Gazzaley et al., 2005), and processing speed (Salthouse et al., 2010). Recent studies have used 1/f slope of the power spectrum as a marker of neural noise and found it to be even predictive of N900 lexical prediction (Dave et al., 2018), measures of working memory cognitive load (Voytek et al., 2020) and in grammar learning (Cross et al., 2020). Considering these results, 1/f noise does not necessarily seem to mediate behaviour in a restricted cognitive domain of interest rather it seems to have much broader repercussion on the overall cognitive processes and brain functions. Therefore, we examined the effect of 1/f noise on cognitive and metacognitive measures during resting-state brain dynamics and correlated those neural measures with multiple aspects of working memory performance through lifespan, instead of focusing on multiple task categories. First, we hypothesised any specific change in the background 1/f slope and offset would index change in the age-associated shift in the baseline neural noise and in turn would impact oscillations in the different frequency bands associated with healthy aging process. Secondly, we hypothesized age-associated 1/f neural differences would correlate to all the behavioural measures suggesting more global changes associated with different aspects e.g., processing speed, cognitive load, accuracy and metacognitive awareness of the participants in a memory task, whereas differences in the oscillations mediated by relevant frequency bands theta, alpha and beta would correspond more to specific behavioural measures reaction time and speed of processing in the same task.

Although behavioural and neuroimaging studies have shown that neural noise increases with age, these studies have relied upon proxies (e.g., reaction time) for neural noise. In this study, we have used a parameterisation model developed by Donoghue et al., 2020, which parametrize the two components and avoid conflating them with one another, where 1/f noise is well characterised by the line in semi-log or log-log of frequency spectra. Dissociating these two aspects of the brain signals naturally give a more accurate estimate of the neural oscillations and normative changes associated with healthy aging and would provide a refined understanding of the aging process in both rest and task conditions and the relationship thereof. Hence, a better characterization of normative features associated with healthy aging would also inadvertently provide a better understanding of how these neural mechanisms affect cognition in general. Most studies have looked at the oscillatory changes while the participant performs a task and correlated the changes with behavioural measures (Moran et al., 2010; need more citation and reference) in an age-stratified manner with a smaller sample of aging participants. One distinct approach from the previous studies here is that we leverage the big data by testing the neural noise hypothesis using a large publicly available dataset by tracking the neural noise dynamics sufficiently at all the important milestones of healthy adult lifespan. By regressing out age, we show that how aperiodic change in 1/f could predict sufficiently well various aspects of memory tasks and has the potential to predict chronological age from task data. In summary, we argue that the normative brain oscillations and dynamical features in the resting-state brain dynamics is a crucial gateway to understand overarching goals of understanding pathological changes. However, the difference in oscillations with aging are often notoriously difficult to understand as they co-exist with underlying change in aperiodic 1/f dynamics (neural noise) and may crucially hinder understanding the patterns associated with cognitive state and aspects of task performance. In this work, we have been able to dissociate those two dynamics and demonstrate how the age associated baseline shift in neuronal dynamics might be responsible for the differences in the task-induced changes across age.

## METHODS

### 1. Participants

The Cambridge Centre for Aging and Neuroscience (Cam-CAN) is a large scale, multimodal, cross-sectional adult life-span (18-88) population-based study. The Cam-CAN consists of 2 stages. In stage 1, 2681 participants had gone through general cognitive assessments at their home. Tests for hearing, vision, balance and speeded response were also assessed. Additionally, measures taken in stage 1 served to screen participants for stage 2. Those with poor hearing, poor vision, with neurological diseases such as stroke, epilepsy or a score less than 25 in MMSE (cognitive assessment examination) were excluded from the further participation. From stage 1 to stage 2, 700 participants were screened (50 men and 50 women from each age band). All screened participants were recruited for testing at the Medical Research Council (UK) Cognition and Brain Sciences Unit (MRC-CBSU) in Cambridge, UK. In this stage, MRI scans, MEG recordings and cognitive task data were collected, all the participants performed a range of psychological tests and neuroimaging assessments, but only the MEG RS data and VSTM task data are included in this study. Out of 700 participants, Magnetoencephalogram (MEG) data from 650 subjects were available. Age values of participants were divided into 4 age groups for categorical analysis (See Methods). Young Adults (YA), Middle Elderly (ME), Middle-Late (ML), Older Adults (OA), Similar grouping has been performed in our previous study also. 70 participants were randomly chosen from each age group resulting in total of N=280 subjects comprise of all four important stages of adult lifespan (Chan et al., 2014; Sahoo et al., 2020).

**TABLE 1:**
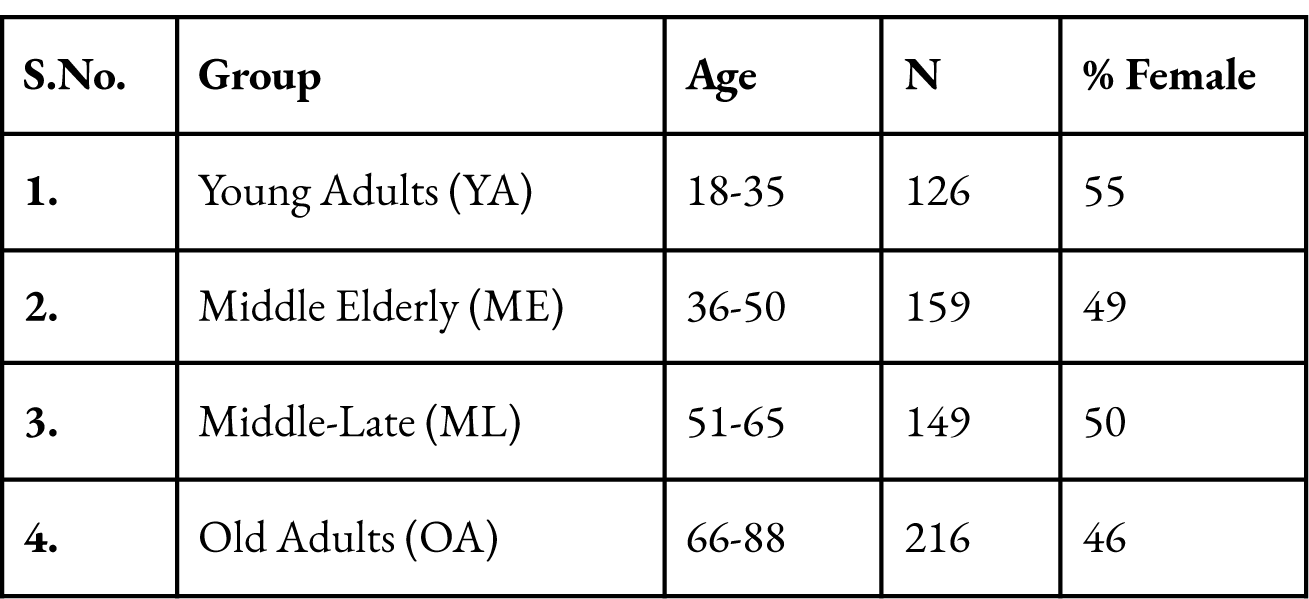
Each representative age group.

### 2. Data acquisition

#### 2.1 MEG Resting-State data

MEG Data used for this study were obtained from the CamCAN repository (available at http://www.mrc-cbu.cam.ac.uk/datasets/camcan/) (Taylor et al., 2007; Shafto et al., 2014). For all the 700 participants, MEG data were collected by Elekta Neuromag, Helsinki at MRC-CBSU using 306 channels, consisting of 102 magnetometers and 204 orthogonal planar gradiometers. MEG data collection was done in a light magnetically shielded room (MSR). A high pass filter of 0.03 Hz cutoff was used to sampled the data at 1000Hz. Head-Position Indicator (HPI) coils were used to continuously assess the head position within the MEG helmet. To monitor blinks and eye-movements, two pairs of bipolar electrodes were used to record horizontal and vertical electrooculogram signals. To monitor pulse-related artefacts, one pair of electrodes were used to record electrocardiogram signals. MEG data collected for resting-state required the participants to sit still for a minimum of 8mins and 40 sec with their eyes closed. From this subset, 280 participants were included in the present study (70 in each group).

#### 2.2 VSTM Stimuli and Task

In CamCAN, the design was adapted from Zhang et al., 2008 (**Figure 1**). On each trial, participants were presented with 1,2,3, or 4 coloured discs (mimicking different memory load conditions) for 250ms. Following that, a blank screen was presented for 900ms to hold those colours in memory. One of the original locations was highlighted by a thick black border (acting as a probe for participants to remember the colour at that location), and at the same time, a response colour wheel was presented. Participants had as much time as required to report by touching or clicking, as accurately as possible the remembered hue of the highlighted disc. No feedback was given. After every trial, 830 ms fixation period was there. Participants indicated their lack of confidence in the precision of the colour (metacognitive awareness) by the length of the time they hold down the finger onto the point. Participants complete two blocks of 112 trials, with memory load (1,2,3 or 4) counterbalanced and randomly intermixed. For each set size (memory load), the following measures were estimated by fitting the error distribution with a mixture model of von-mises and uniform distributions, proposed by Zang & luck (2008) and modified by Bays and Husain (2008). For detailed analysis refer to Zhang et al., 2008; Mitchell et al., (2018).

**Figure 1.**
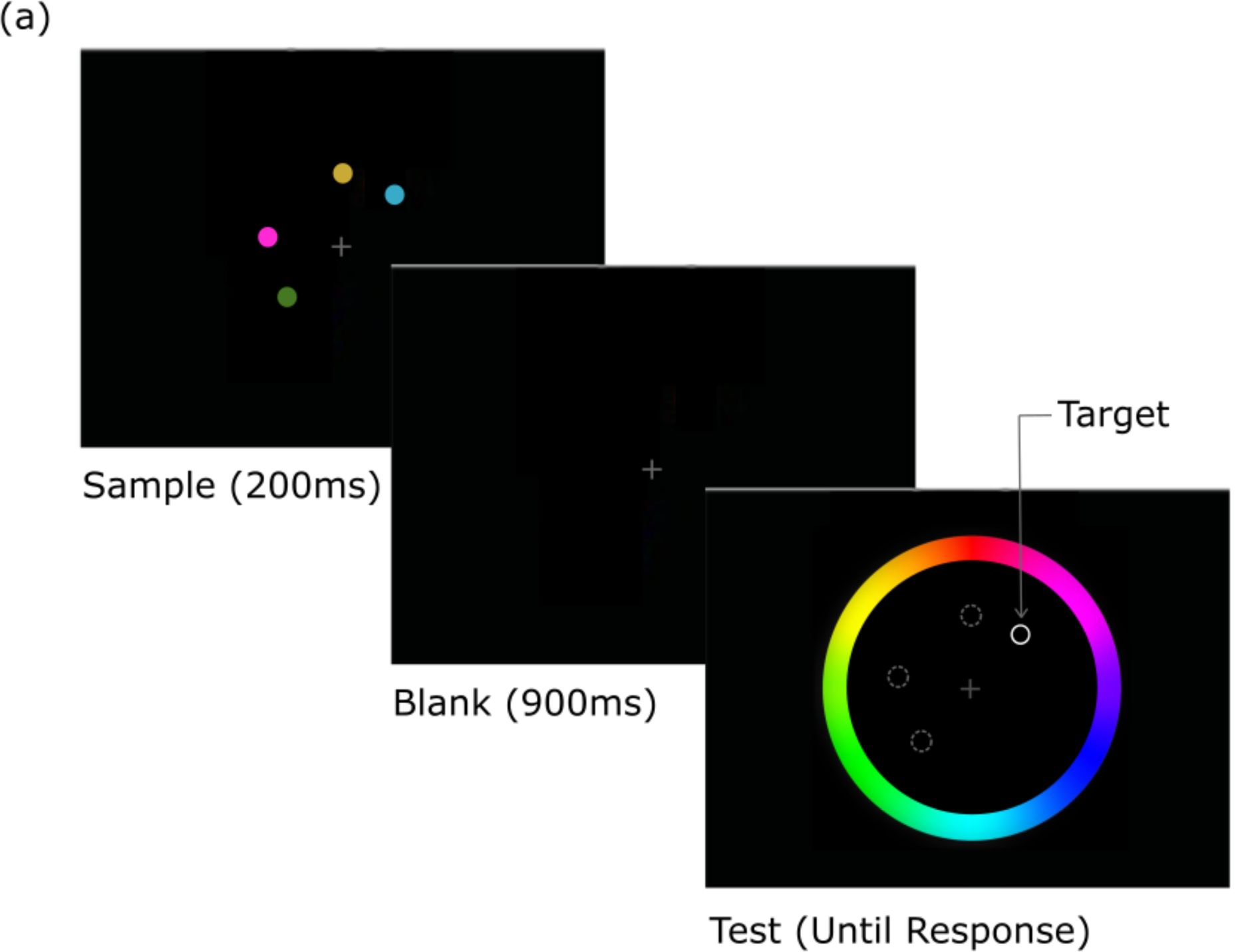
Experimental design of the colour recalltask. Example trial, with memory load of 4 items. (Data were taken from CamCAN repository; Adapted from Mitchell et al., (2018)).

**TABLE 2:** Estimated measures of VSTM task.

| S.No | Variable | Description |
| --- | --- | --- |
| 1. | **Precision** | Accuracy of reportable items (reciprocal of Std. deviation of fitted von-mises) (degrees) |
| 2. | **RT** | Median Reaction time (ms) |
| 3. | **K(VSTM Capacity)** | Number of reportable items (k-score) |
| 4. | **Mean Uncertainty** | Size of confidence interval within which answer is thought to lie (degrees) |

#### 2.3 MEG Data Preprocessing

MEG processed data was provided by Cam-CAN. Preprocessing pipeline included temporal signal space separation, applied on continuous MEG data to remove noise from the HPI coils, environmental sources and continuous head motion correction. For removing the main frequency noise (50 Hz notch filter) and to reconstruct any noisy channel, max filter was used. More details about data acquisition and pre-processing have been presented elsewhere (Shafto et al., 2014; Taylor et al., 2007). Additionally, we performed independent component analysis (ICA) to get rid of the artifacts and harmonics in the signal (Sahoo et al., 2020).

#### 2.4 Data Analysis

All data were analyzed in MatLab and python using custom scripts. Python MNE for preprocessing, standard python libraries including *Scipy, Pandas* and *NumPy* for data management and processing, python-*matplotlib* and *seaborn* for data visualisation were used in this study. The analysis pipeline is concisely represented in **Figure 2**.

**Figure 2.**
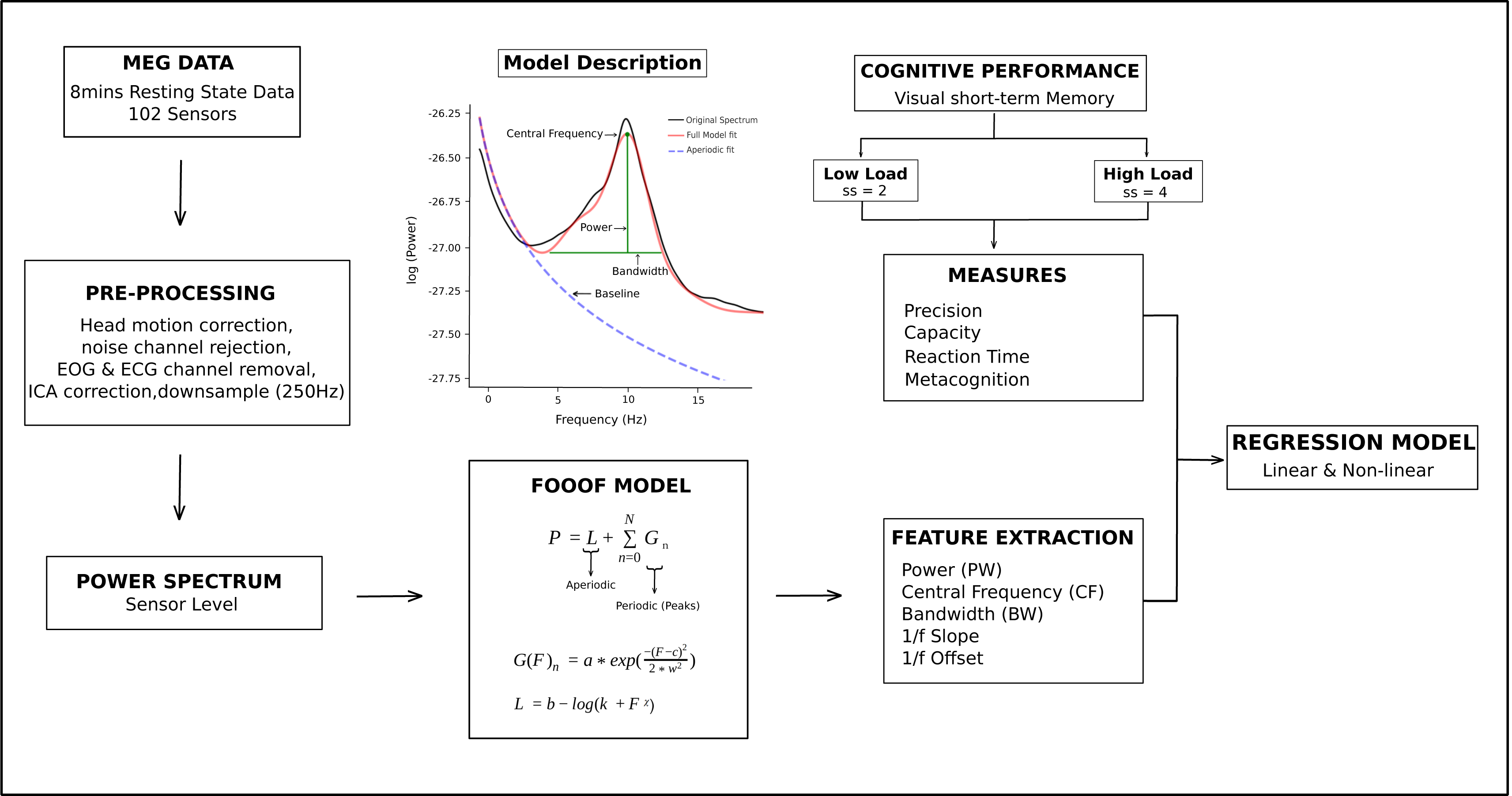
Data processing and analyses pipeline.

##### 2.4.1 Power Spectral Density (PSD) using Welch’s periodogram method

The Power spectrum *s*_*xx*_(*f*) of a signal *x*(*t*) capture how the strength of the signal is distributed in the frequency domain. Using Fast Fourier Transform (FFT), (a variant of Fourier Analysis), the representation of raw signal (time or space) is transformed into a frequency representation of the signal.

Processed MEG data provided in ‘.fif’ format was analysed using Fieldtrip toolbox (Oostenveld et al., 2011). Data for each N=280 subjects were first downsampled from 1000 Hz to 250Hz. The frequency resolution was held at 0.05 Hz. Power spectral density (PSD) was estimated using Welch’s periodogram method implemented in MATLAB 2019b, which has 2 additional steps before computing the FT. First, the estimation divides the signal into *n* segments (creating epochs) with some overlap. Subsequently, the epoched signal further windowed using a window function.

For each participant, 102 magnetometer sensor’s time series data resulted in a matrix of size 102 *X T* where *T* correspond to the number of time points. Each sensor’s *x*_c(t)_ time series *x*_c(t)_ was further divided into segments of 20s without any overlap. For each segment, the spectrum was calculated and then was averaged. For estimation of global spectrum, representative of each subject i.e., *S*_I(*f*)_, the grand average across the spectrum of all the magnetometer sensors are provided below

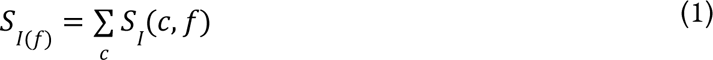

For each participant, resulted power spectrum matrix was *v X c*. For group-wise analysis, each participant’s spectrum was averaged across sensors of interest.

##### 2.4.2 Extracting Periodic and Aperiodic Features using a Parameterization model

To detangle the periodic (oscillatory) component from the aperiodic properties of the signal power spectra. we used a recently proposed parameterization model, fitting-oscillations-and-one-over-f (FOOOF toolbox) (for a full description refer to Donoghue et al., 2020). In brief, the PSDs calculated using *pwelch* was given as an input to the model, which considers PSDs as a linear sum of aperiodic 1/f like characteristics of neural power spectra and it is entirely described by the aperiodic “exponent” and “offset”. Periodic components describe putative oscillations that describe power above aperiodic component (so-called ‘peaks’, simulated as Gaussian function; are described by Peak Frequency in hertz (Hz); peak power over and above the 1/f signal in arbitrary units (au) and bandwidth which describes the spread also measured in the unit of Hz). The simulation, for a power spectrum *P* is described as follows,

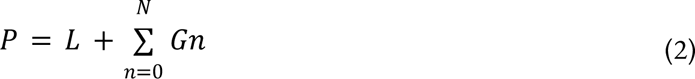

Where P is the linear sum of the aperiodic signal *‘L’* and N Gaussian peaks *‘Gn’.* For each peak, Gaussian function ‘*G*_n_’ is fitted which is modelled as:

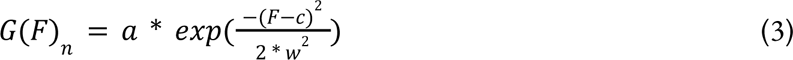

Where ‘a’ denotes the amplitude, *‘c’* denotes the central frequency, *‘w’* denotes the bandwidth of the Gaussian. ‘F’ is the frequency vector. Subsequently, all fitted Gaussians were subtracted from the original power spectrum to get a peak-removed power spectrum (PRPS). Finally, a 1/f signal is estimated from this PRPS using Eq. (4), representing the actual cortical noise. Exponential function in semilog-power space (logged power values and linear frequencies) is used to model the aperiodic signal (initial and final fit both), ‘L’, as:

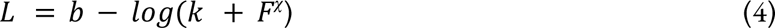

Where *‘b’* denotes the broadband offset, *‘χ’* is the slope, and *‘k’* is the knee parameter, which depends on the bend in the aperiodic signal. The FOOOF model was fitted across the frequency range of 1 to 45 Hz in fixed mode, as no knee was expected in the MEG recordings across 1-50 Hz frequency range (Miller et al., 2009) Algorithm was implemented using custom python scripts on the python3 version.

The Model was fit per subject and output parameters were averaged across subjects for each group **(Figure. 3(A))**. The settings for the algorithm were set as: (1) peak_width_limits = [0.5, 12]; (2) min_peak_height = 0; (3) max_n_peaks = 12; (4) peak_threshold = 2; (5) aperiodic_mode = “fixed”; and (6) verbos = ‘True’.

Oscillations were pot-hoc grouped into theta (θ, 4-8 Hz), alpha (8-12 Hz), and Beta (β, 13-30 Hz). For estimating the topographical dynamical changes, the brain was segmented into 5 non-overlapping regions: frontal (number of sensors = 26), parietal (number of sensors = 26), occipital (number of sensors =24), right and left temporal (number of sensors = 26).

##### 2.5.3 Band Ratio Measures

Additionally, we estimated the band ratios which reflect the quantitative measure of oscillatory activity and are investigated in different cognitive processes; however, they also get impacted by the 1/f background noise (Donoghue et al., 2020). After removing the aperiodic signal using a parametrization method proposed by the Fooof toolbox, periodic values were estimated. Thus after implementing appropriate parametrization of the aperiodic component of the signal power spectra band ratio values were re-estimated to indicate the true power changes and finally, were grouped into different frequency bands of interest. For each participant, we calculated the ratio of periodic components of different frequencies and averaged across participants for age bin-wise distribution. Band ratio of all the periodic components for each frequency band was then calculated by dividing the average of low band periodic features by the average of high band periodic features. We calculated frequency-specific band ratios of all periodic features.

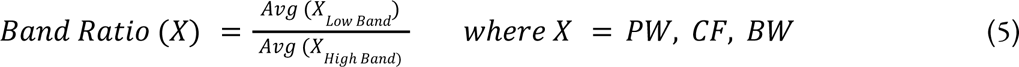

##### 2.5.4 Statistical Analysis

We performed both categorical as well as continuous analysis to capture different aspects of age-associated functional differences. For the continuous analysis, we divided the total number of participants into bins of 5 years starting from 18 years, a total of 14 bins and the centre value was taken to be the representative age for each bin. For the categorical analysis, we divided data into the following age stratifications (18-35 years, 36-50 years, 51-64 years, 66-88 years) to get insights about different important stages of adult lifespan and comparison with previous works (Chan et al., 2014; Sahoo et al., 2020).

###### Correlation Analysis

Depending on the data distribution, Pearson or Spearman’s correlation was used to estimate the strength between two variables. Estimated aforementioned functional changes (oscillatory, aperiodic, band-ratio measures) and VSTM task measures were correlated with age. Finally, VSTM task measures were then correlated with those functional changes.

###### Regression Analysis

Linear and Non-linear Regression were performed separately considering each Power, central frequency, Bandwidth, slope, offset and band-ratios of periodic features, as the estimated measures (R) of functional changes and Precision, Reaction Time (RT), Metacognitive awareness (d) and memory capacity (k), as the estimated measures (R) of the VSTM behavioural task, while keeping age as an explanatory variable.

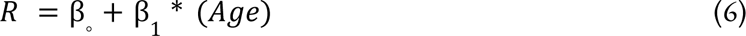

Linear regression was performed using *fitlm* matlab function. To capture the potential non-linear effects of age, we also added 2^nd^ order polynomial terms to the model, such as:

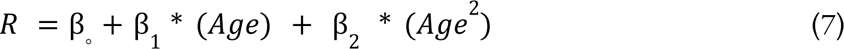

Linear regression was also performed considering each VSTM task measures as a response variable (R) and the functional measures as the explanatory variable (E). (Detailed report is provided in supplementary)

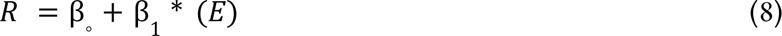

All regression tables are provided in the supplementary document. For estimating the significance, first normality of the data distribution was assessed using the Kolmogorov Smirnov test. Based on the data distribution, parametric (t-test) or nonparametric (Wilcoxon rank-sum test) was performed.

## RESULTS

From the parametrization model fit, all the simulated Gaussian peaks were removed to analyse the background signal. Thereafter, the aperiodic component of the signal was fitted in the log-log space line (**Extended fig. 3-2**) from which 1/f Slope and offset were extracted from the Fooof model for each individual participant. Periodic features Central Frequency (CF), Power (PW), Band Width (BW) were estimated using peak parameters from the fitted model (refer to the materials and methods section). To check if the parameterisation using the simulated fooof model is able to capture lifespan associated changes, we first simulated the Fooof model for young and old adults. The model well captured the well-established lifespan associated slowing down of Peak alpha frequency (PAF)(**Figure 3**). Original spectrum, aperiodic fit and full model are being depicted in **Figure 3(B)(C)** for YA and OA group respectively (refer **Extended fig. 3-1** for model’s output parameters of ME and ML groups).

**Figure 3.**
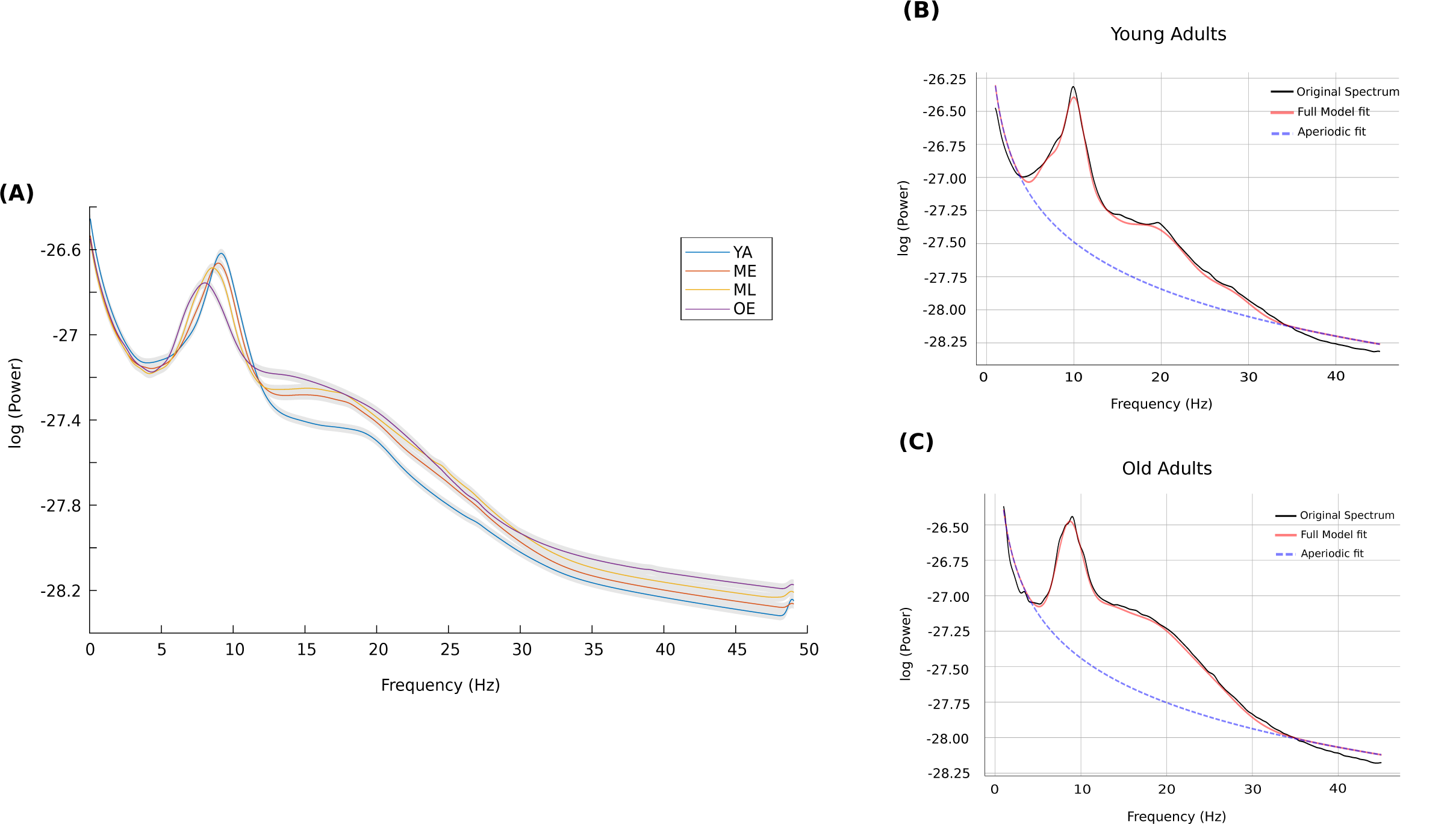
Parameterization using FOOOF Model. **(A)** Power Spectrum of all age-groups after removing the 1/f aperiodic component **(B) & (C)** FOOOF Model fit for Young and Old Adults.

To capture the dynamical changes in the dominant oscillations (highest power peak across all frequencies) across the adult lifespan, the central frequency, power and bandwidth of the dominant oscillations were also extracted for young and old adults. No significant difference was found in the frequencies of dominant oscillation however, the power of the respective dominant frequencies was found to be significantly different between YA and OA (**Extended fig. 5-1**).

**Figure 5.**
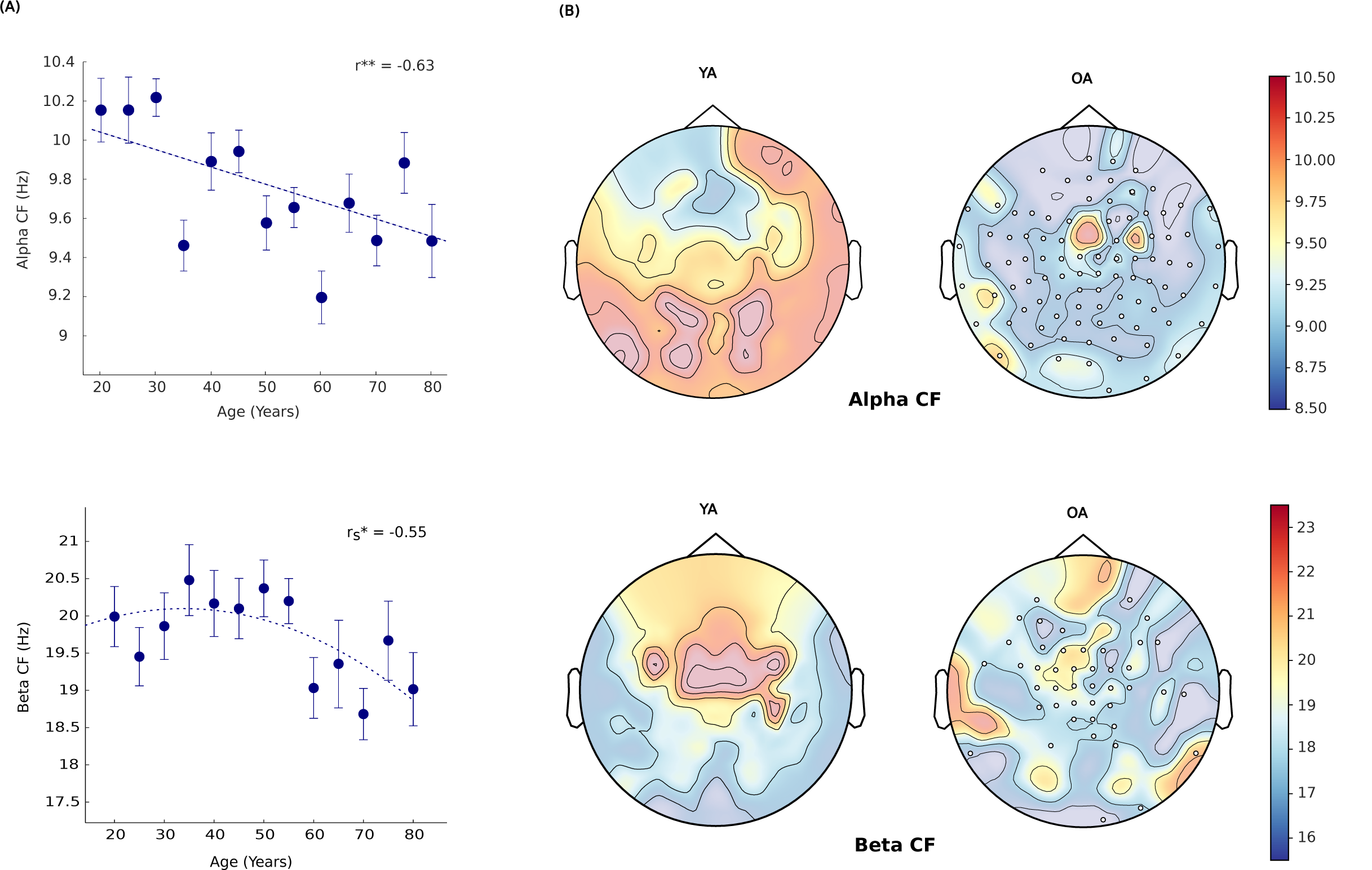
Alpha and Beta peak frequency as a f unction of Age (A) Top: PAF as a function of age. Bottom: Beta peak frequency with age. ‘r’ represents the correlation coefficient. The dashed line represents a linear regression fit. Error bar denotes SEM. **(B)** Top: Spatial Topography for PAF and Beta peak frequency for Young (YA) and Old adults (OA). Clusters of sensors with negative differences which contribute to the decrease are represented as white dots.

### 1. Topographical distribution of Aperiodic componentof the signal with Age

#### Increase in Aperiodic 1/f slope and decrease in 1/fintercept

We found that aperiodic 1/f slope increases significantly (β_1_ =+0.0034901, R^2^ = 0.584, p = 0.003) whereas 1/f offset doesn’t show significant decrease across the adult lifespan (β_1_ = -0.0033423, R^2^ = 0.3, p =0.03) (**Figure 4(A), (B)**, **Extended fig. 3-3**). Categorical analysis also confirmed significant difference in the 1/f slope between the OA vs YA (t(140) = 4.38, p <0.0001), ML vs YA (t(140) = 4.07, p = 0.02), ME vs YA (t(140) = 2.7749, p = 0.007), ME vs OA (t(140) = -2.4581, p = 0.02). Categorical difference in 1/f offset was also found between OA vs YA (t(140) = 2.0345, p = 0.0457) and ML vs YA (t(140) = -2.3441, p = 0.02) (**Figure 4(A), 4(B)**). Within group analysis revealed more variability in aperiodic features in the older adults (Slope: SEM = 0.023; Offset: SEM = 0.0404) compared to young adults (Slope: SEM = 0.014; Offset: SEM = 0.0364) (**Extended figures 3-4, 2-4**). **Figure 4C** and **Figure 4D** shows variability in spatial topographies of aperiodic 1/f slope and offset for Young and Old adults.

**Figure 4.**
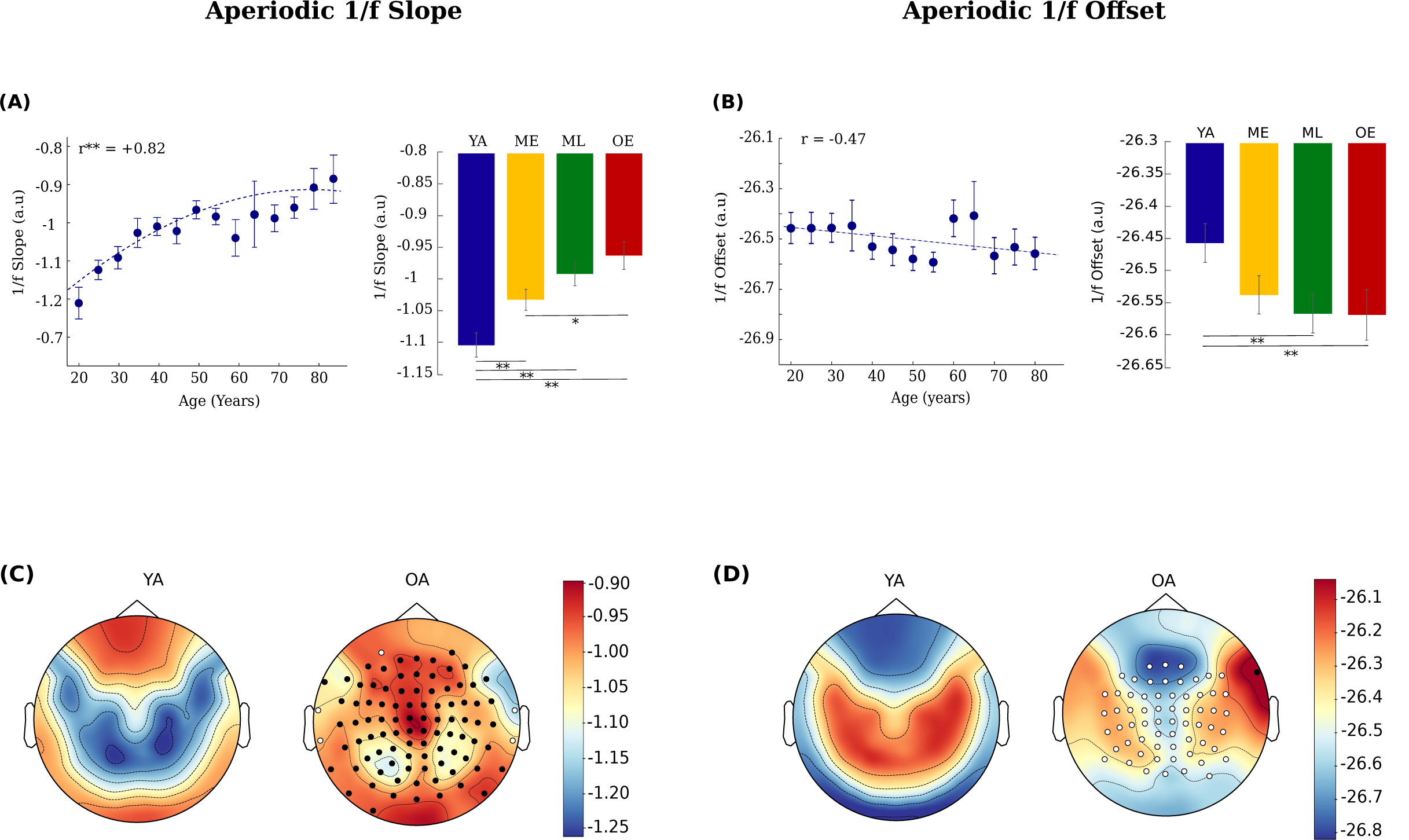
Aperiodic 1/f Slope and Offset. **(A)** Left: 1/f slope as a function of age. Right: 1/f slope for four age groups. **(B)** Left: 1/f Offset as a function of age. Right: 1/f Offset for four age groups. ‘r’ represents the correlation value. The dashed line represents a linear regression fit. Error bar denotes SEM. **(C)(D)** Aperiodic 1/f slope and 1/f offset spatial topography for Young (YA) and Old adults (OA). Clusters of sensors with significant positive and negative differences in 1/f slope and 1/f offset between the OA and YA group are represented with black and white dots, respectively.

### 2. Topographical distribution of oscillatory component with age

#### Age-associated slowing of Central Alpha frequencyand Beta frequency

For each participant, PF was quantified by estimating the peak power value within the 8-12 Hz and 13-30 Hz for alpha and beta range, respectively. Each participant’s PF was then averaged to get the group-wise estimation of Central Alpha Frequency (CAF). Visual inspection revealed bin 65 to be the outlier (for CAF). After removing the outlier, significant linear age-related decline was found (β_1_ = -0.010234, R^2^= 0.4, p = 0.02) however, central beta frequency (CBF) showed non-linear decrease with age (β_1_ = -0.024068, R^2^= 0.462, p = 0.007) (**Figure 5(A)**). Categorical analysis also revealed significant CAF differences between YA vs OA (t(140) = 4.7551 p =0.00001), YA vs ME (t(140) = 3.4198, p = 0.001) and YA vs ML (t(1400 = 4.8826, p = 0.000001), and for CBF between YA vs OA (t(140) = 1.912, p = 0.03). Almost all sensors were found to be contributing to the decrease in CAF in OA whereas the decrease in CBF was mainly contributed by the central sensors (**Figure 5(B)**).

#### Functional Power change in Alpha, Theta and Beta frequency with age

We found a robust decline of Alpha power with age (β_1_ = -0.0059263, R^2^= 0.75, p =0.00005) (**Figure 6(A)**). Visual inspection suggests that sensor level Alpha power difference was mainly contributed by the occipital, parietal and left temporal sensors (**Figure 6(B)**). Significant difference was found between OA vs YA (t(140) = -3.038, p = 0.003), OA vs ME (t(140) = -2.2008, p = 0.03) and OA vs ML (t(140) = -2.2252, p = 0.029). Older adults showed higher Theta power (M = 0.56 ± 0.04) than younger adults (M= 0.32 ± 0.02) (t(140) = 2.4733, p = 0.023) . Significant age effect was also observed with increase in theta power (β_1_ = 0.0050947, R^2^= 0.363, p = 0.022) (**Figure 6(A)**), which was mainly contributed by the temporal sensors. In addition, aging was also associated with an increase in Beta Power (β_1_ = 0.002496, R^2^= 0.70, p = 0.0001) (**Figure 6(A)**). Spatial topographies showed Central and frontal sensors to be contributing to this age-related increase in global beta power (**Figure 6(B)**). Categorical analysis revealed significant differences in Beta power between the YA vs OA (t(140) = -4.3693, p = 0.00004), YA vs ME (t(140) = -3.0103, p < 0.003), and YA vs ML (t(140) = -4.4158, p = 0.00003). **Extended Fig. 6-1** shows the sensor-wise distribution of frequency specific power as a function of age.

**Figure 6.**
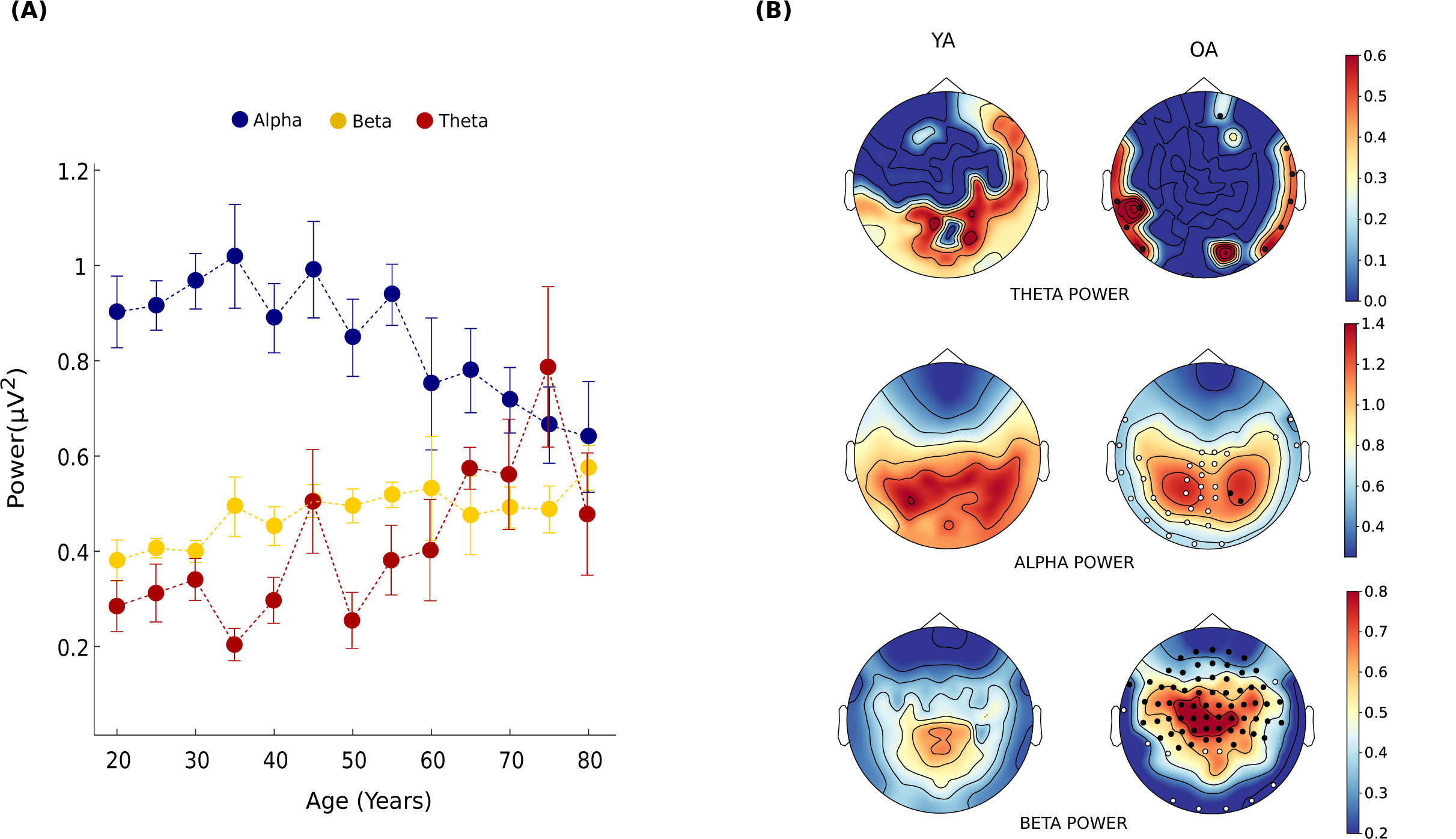
Parameterised Global Power as a Function of Age. **(A)** Increase in Theta and Beta power whereas a decrease in Alpha power with Age. Errorbar represents SEM. **(B)** Spatial Power topography of Theta, Alpha and Beta for young (YA) and old adults (OA). Clusters of sensors with significant positive and negative differences between the OA and YA group are represented with black and white dots, respectively.

#### Increase in Beta bandwidth with age

Bandwidth reflects the spread of power in the respective frequency range, which for the Beta band was found to be increased across the adult lifespan (β_1_ = 0.040345, R^2^= 0.58 p = 0.001) (**Figure 7(A)**). Significant group-wise difference was also seen between YA vs OA (t(140) = -3.1586, p = 0.0024), YA vs ME(t(140) = -1.9843, p = 0.049), suggesting that as we age, the Beta power tends to spread more across frequency range. This increase was mainly observed over left temporal and central sensors (**Figure 7(B)**). Bandwidth for Alpha and Theta frequency band did not differ across age groups (**Extended fig. 7-1**). For sensor topography refer to **Extended fig. 7-2**.

**Figure 7.**
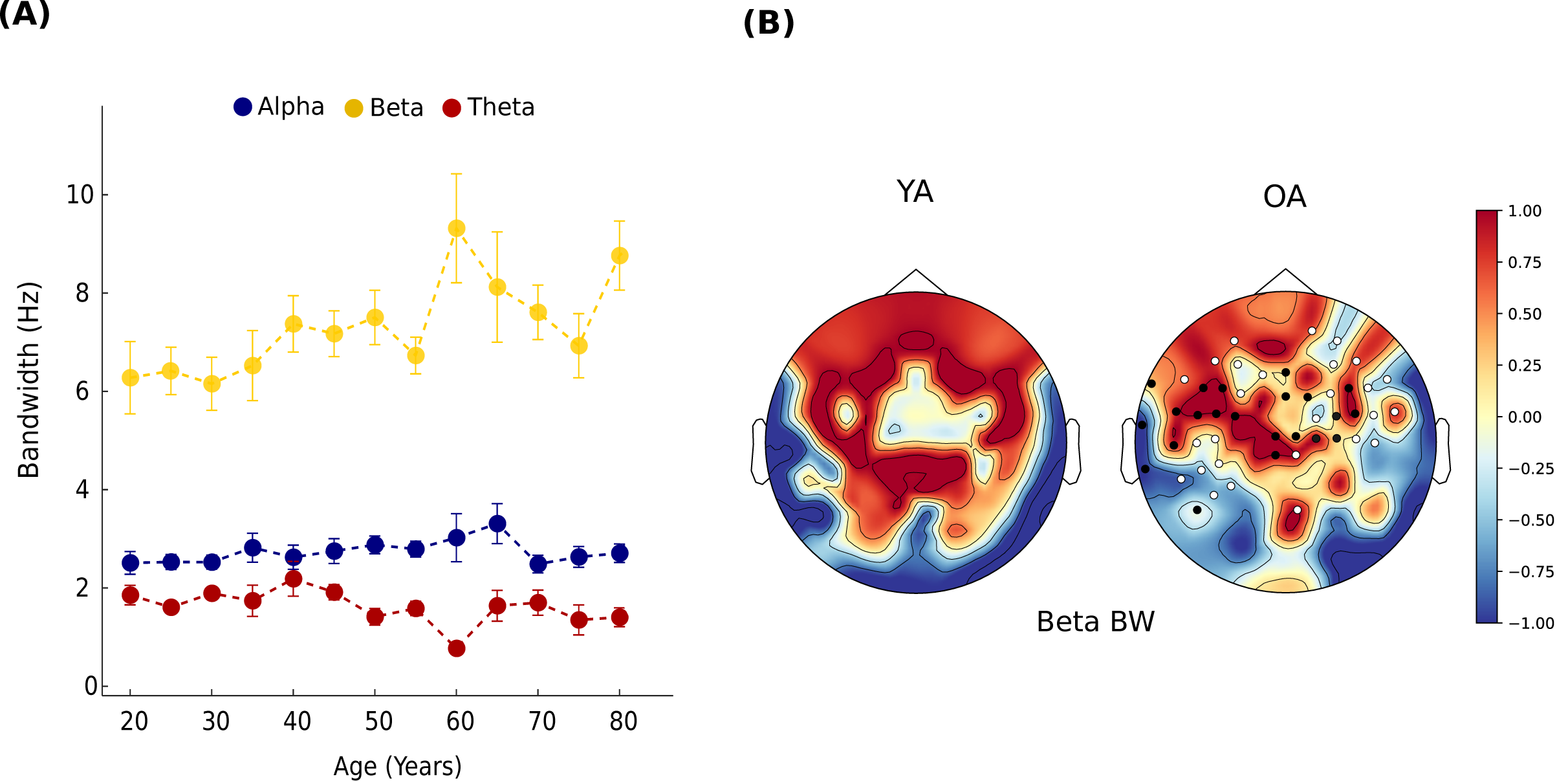
Global frequency-specific Bandwidth with Age. **(A)** Bar graph for each age group, representing bandwidth for each frequency band. **(B)** Spatial topography of Beta BW for young (YA) and old adults (OA). Clusters of sensors with significant positive and negative differences between the OA and YA group are represented with black and white dots, respectively.

### 3. Band-Ratios measures of periodic features withage

Band ratio measures have been argued to be a marker of various cognitive measures in healthy adults as well as in pathological conditions (Trammell et al., 2017; Kamiński et al., 2011; Schutter et al., 2005) which also get affected by 1/f noise. We investigated how these Global band ratios change with age after effectively removing the background 1/f noise. We looked at Theta/Alpha (*θ*/*α*), Theta/Beta (*θ*/*β*) and Alpha/Beta (*α*/*β*) Band ratios, where the ratio of all periodic features (PW, CF, BW) was analyzed for each frequency band. For all band ratio measures, we calculated correlations between the spectral features of each oscillation-band and age. Here we showed the global change (averaged across all sensors) in the band ratio measures across the lifespan.

For the Central frequency ratio, we found *α*/*β* ratio to vary non-linearly (quadratic) with age (*β*_1_ = -0.0059138, R^2^=0.61 p = 0.005), whereby first decreases for middle age and subsequently an increase for older age participants suggesting an overall U-shaped response of *α*/*β* ratio through lifespan (**Figure 8(B)**). This age-associated nonlinear change was mostly observed in frontal and parietal sensors (**Figure 8(A)**). For age categories, we found a significant difference between OA vs ME (t(134) = 2.38, p = 0.018), OA vs ML (t(134) = 3.19, p = 0.0018), YA vs ME (t(138) = 3.30, p = 0.0012) and YA vs ML (t(138) = 4.09, p = 0.00007). No significant difference was found between the categorical age groups for *θ*/*α* and *θ*/*β* peak ratios.

**Figure 8.**
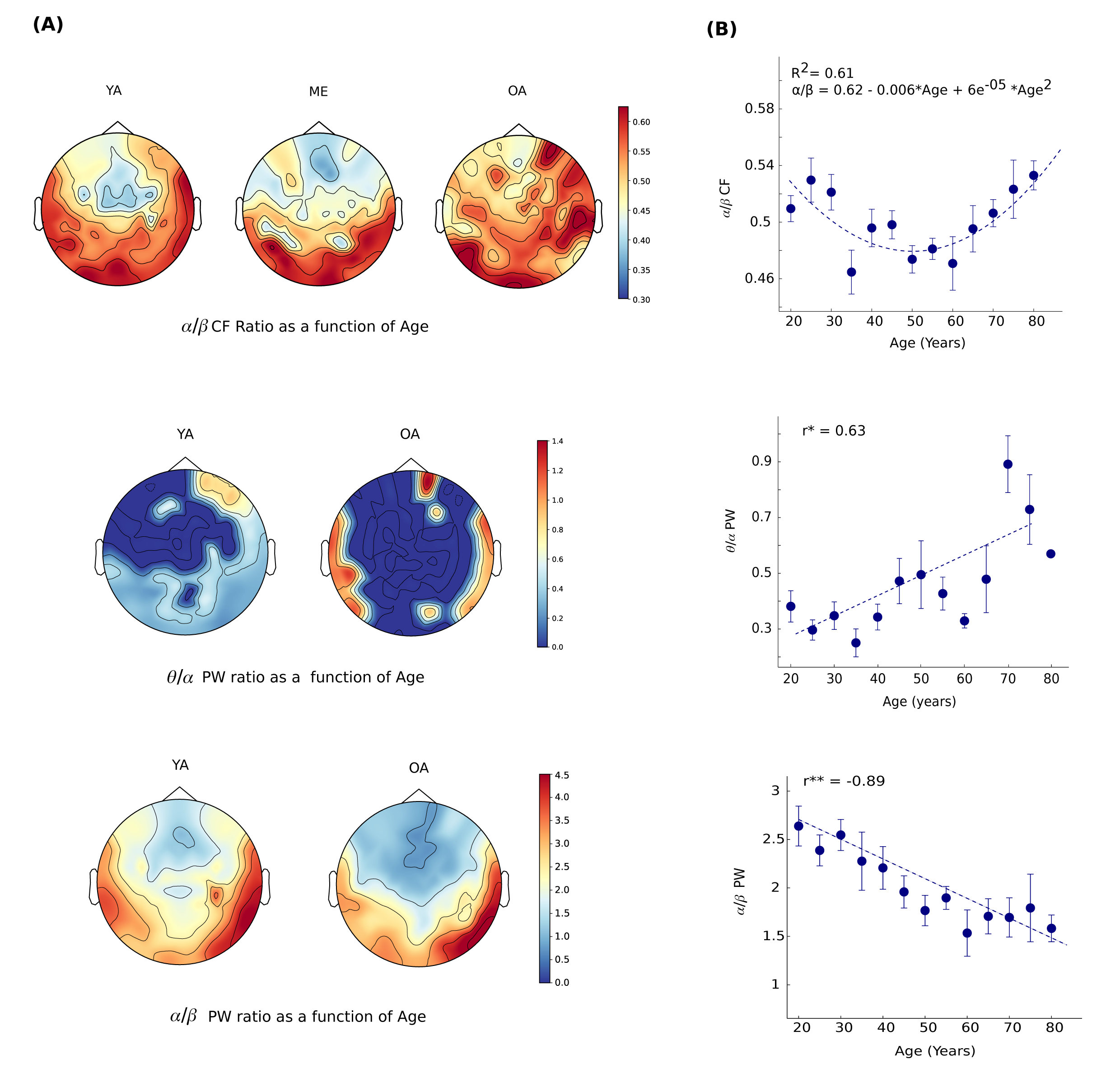
Spatial topography of band-ratio measuresas a f unction of age. (A) Spatial topography of Alpha/Beta peak frequency ratio (*α*/*β* CF) (top), Alpha/Beta power ratio (*α*/*β* PW) (centre) and Theta/Alpha power ratio (*θ*/*α*) (bottom) for young (YA) and old adults (OA). (B) Regression fit model for each of the aforementioned ratio measures keeping age as an explanatory variable. Error bar represents SEM. R^2^ represents goodness of fit and ‘r’ represents the correlation coefficient.

Power Ratio of *θ*/*α* was found to be positively correlated with age (*β*_1_ = 0.0057613, R^2^= 0.40, p = 0.02) whereas *α*/*β* power ratio was negatively correlated with age (*β*_1_ = -0.018116, R^2^= 0.85, p = 0.000001) (**Figure 8(B)**). Significant Categorical difference was found for *θ*/*α* power ratio between YA vs OA (t(136) = 4.9615, p = 0.000002), YA vs ME (t(138) = 2.75, p = 0.0067), YA vs ML (t(138) = 4.92, p = 0.000002), ME vs OA (t(134) = 2.24, p = 0.02) No significant correlation was found for *θ*/*β* power ratios with age (R^2^= 0.2, p = 0.1). For *α*/*β* power ratio, significant categorical difference was found between YA vs OA (t(76) = -4.6, p = 0.00001), ME vs OA (t(59) = -3.33, p = 0.0015), and ML vs OA (t(62) = -2.46, p = 0.01). No significant difference was found between the categorical age groups for *θ*/*β* power ratio.

Bandwidth ratio of *θ*/*β* and *θ*/*α* was found to be negatively correlated with age (**Extended fig. 8-3,4**). Categorical analysis revealed differences between the YA vs OA (t(80) = 2.21, p = 0.029) for *θ*/*β* bandwidth ratio. No significant difference was found between the categorical age group for *α*/*β* bandwidth ratio.

**Table 3:**
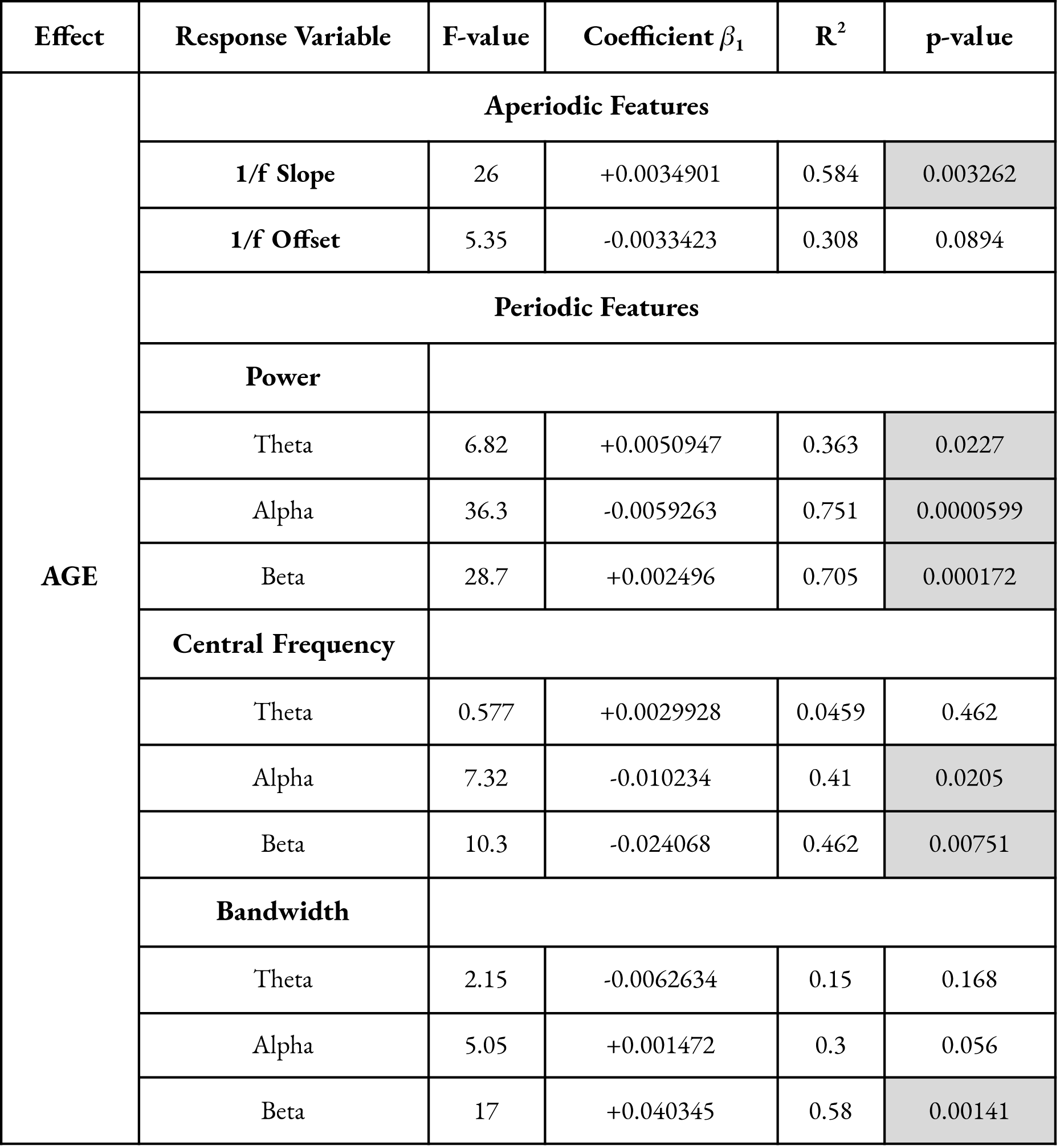
Effect of Age on Periodic and Aperiodic Features.

After getting the normative pattern of true oscillatory changes across age, finally, we tested our hypothesis by carrying out regression analysis whereby keeping 1/f noise, periodic features as an explanatory variable and behavioural measures as response variable (see methods). A detailed Regression table is provided in the supplementary data. All correlations were performed after regressing the ag.

We first analysed the behavioural responses of the same participants in the visual short term memory task to replicate the well-established cognitive decline with age. Grouping of participants in the age groups and bins were done similarly.

### 4. Behavioral Results: Age-Related Cognitive declinereflected in Performance

#### Precision

As expected Precision becomes worse with memory load and age. Overall Precision was high for the set size 1 (61.1% SEM 2%) as compared to set size 2 (48.7% SEM 1.9%), 3 (39.6% SEM 1%) and 4 (39% SEM 0.7%). Continuous analysis revealed significant decrease in precision with age in both low and high load conditions (Low load, r = -0.85, p <0.01, High load, r = -0.61, p < 0.05) **(Figure 9 (A), Extended fig. 9-1**). Categorical analysis between the groups revealed significant differences in the mean of YA vs OA (YA = 0.48 ± 0.008, OA = 0.30 ± 0.005, p < 0.0001), YA vs ME (YA = 0.48 ± 0.008, ME = 0.45 ± 0.007, p < 0.001), YA vs ML (YA = 0.48 ± 0.008, ML = 0.43 ± 0.007, p < 0.0001), and ME vs OA (ME = 0.45 ± 0.007, OA = 0.30 ± 0.005, p < 0.0001) groups. Within group analysis also revealed significant increase in Precision with increase in memory load (**Extended figure 9-2**)

**Figure 9.**
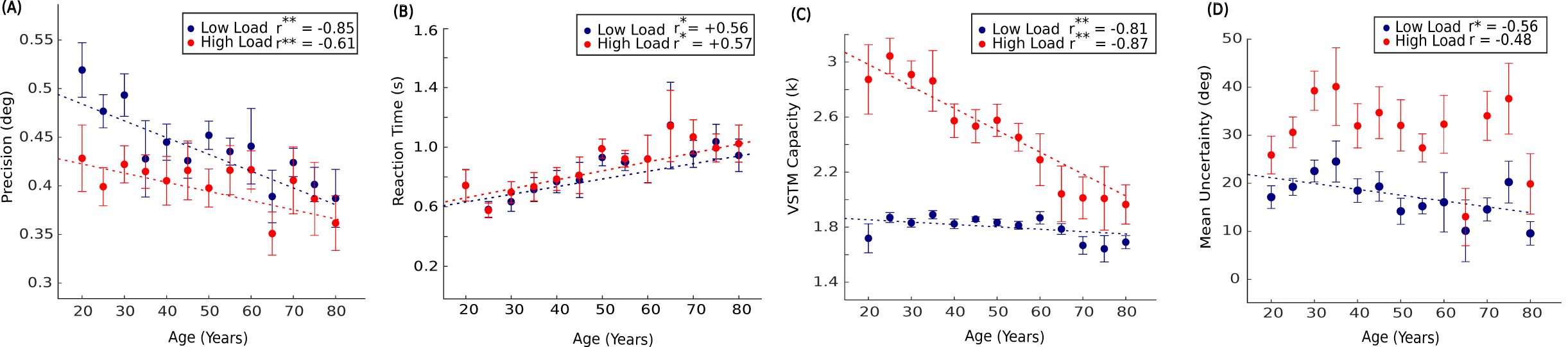
Effect of Memory load and Age on VSTM. VSTM measure **(A)** Precision **(B)** Reaction time **(C)** VSTM capacity (k) **(D)** Mean Uncertainty as a function of age. Low load and high load indicate set size 2 and 4 respectively. The dashed line represents the linear regression fit. Error bar represents the SEM for each age bin. Asterisks indicate significance.

#### VSTM capacity (k)

VSTM capacity was found to decrease with age (Low load (r) = -0.81 p<0.001, High load (r) = -0.87, p <0.001) (**Figure 9(B)**). Categorical analysis between the group revealed significant difference between YA vs OA (YA = 1.84 ± 0.01, OA = 1.66 ± 0.02, p < 0.0001), YA vs ML (YA = 1.84 ± 0.01, ML = 1.79 ± 0.01, p = 0.004), ME vs OA (ME = 1.83 ± 0.01, OA = 1.66 ± 0.02, p < 0.0001), and ML vs OA (ML = 1.79 ± 0.01, OA = 1.66 ± 0.02, p < 0.001).

#### Mean Uncertainty

Subjective Uncertainty was higher in set 4 (29.7 ± 1) as compared to set size 1 (11.8 ± 0.39), 2 (15.6 ± 0.5) and 3 (20.7 ± 0.67). After performing regression and correlation analysis, we found that subjective uncertainty significantly decreases with age in low load condition (Low load (r) = -0.56, p <0.05) (**Figure 9(C)**). Suggesting that Older adults tend to be more confident about their erroneous answers when the load is less. Categorical analysis revealed significant differences in the mean of YA vs OA (YA = 18.5 ± 0.97, OA = 14 ± 1, p < 0.001), YA vs ME (YA = 18.5 ± 0.97, ME = 16.07 ± 0.95, p = 0.02), YA vs ML (YA = 18.5 ± 0.97, ML = 14.85 ± 1.05, p < 0.001), and ME vs OA (ME = 16.07 ± 0.95, OA = 14 ± 1, p = 0.003). Within group analysis also revealed significant increase in subjective uncertainty with increase in memory load (**Extended figure 9-3**)

#### Reaction Time

Overall Reaction time was higher for the set size 4 (910.2 ± 21.6 ms) as compared to set size 1 (878.7 ± 19.9 ms), 2 (870.4 ± 19.9 ms) and 3 (882.6 ± 21.4 ms) but increases significantly with age (Low load (r) = +0.57, p <0.05, High load (r) = +0.56, p <0.05) (**Figure 9(D)**). Group analysis also revealed significant difference between YA vs OA (YA = 668 ± 35.4, OA = 1009 ± 38.9, p < 0.00001), YA vs ME (YA = 668 ± 35.4, ME = 828.8 ± 41.5, p = 0.002), YA vs ML (YA = 668 ± 35.4, ML = 886 ± 33.4, p < 0.0001), ME vs ML (ME = 828.8 ± 41.5, ML = 886 ± 33.4, p = 0.03), ME vs OA (ME = 828.8 ± 41.5, OA = 1009 ± 38.9, p < 0.001), ML vs OA (ML = 886 ± 33.4, OA = 1009 ± 38.9, p = 0.05).

### 5. Aperiodic 1/f slope: Predictive of all measuresof VSTM

We then assessed whether the VSTM performance was impacted by 1/f slope. As hypothesised, RS Aperiodic 1/f noise was found to be predictive of decreased precision (Low Load: r = -0.74, p = 0.002, High Load: r = -0.48, p = 0.08), memory capacity (Low Load: r = -0.68, p = 0.0007, High Load: r =+0.82, p = 0.0003), Mean Uncertainty (Low load: r = -0.58, p =0.03, High Load: r = -0.6, p = 0.02) and increased Reaction Time (Low load: r = +0.56, p = 0.00005, High load: r = +0.57, p = 0.00005) in Visual Short term memory task (**Figure 10)**. However, we didn’t find any correlation between 1/f offset and behavioural measures. As aperiodic 1/f noise mediated a global effect on the VSTM performance, we further wanted to investigate how different oscillatory components mediate changes in the specific behaviour measures in VSTM performance through lifespan.

**Figure 10.**
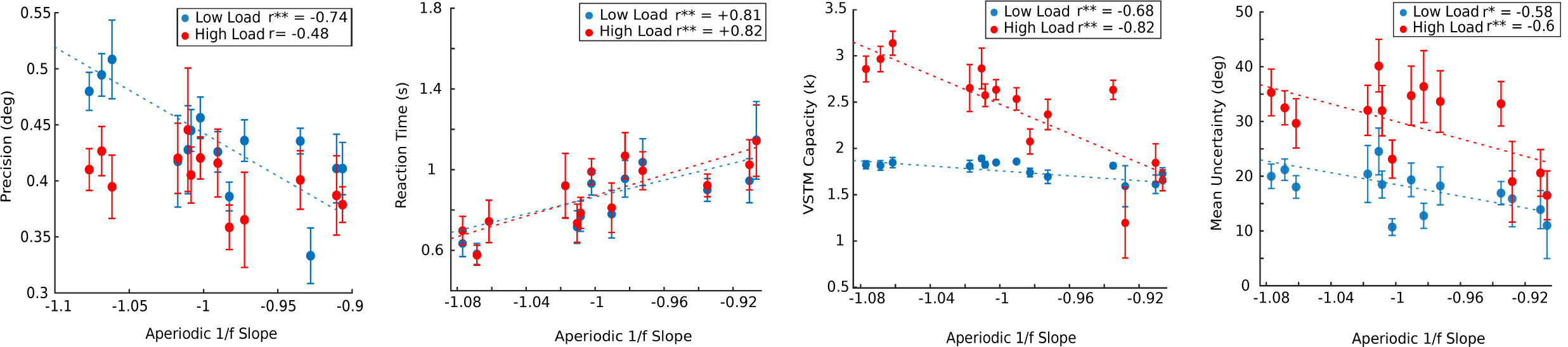
Aperiodic 1/f slope mediating VSTM performance. Linear regression model for VSTM measure **(A)** Precision **(B)** Reaction time **(C)** VSTM capacity (k) **(D)** Mean uncertainty as a response variable and aperiodic 1/f slope as an explanatory variable, after regressing out the age effect. The dashed line represents linear regression fir. Error bar represents SEM. Low load and high load indicates set size 2 and 4 respectively. ‘r’ is Pearson’s coefficient.

### 6. Precision increases with increase in Alpha power and ***α***/***β*** power ratio

Precision was found to be positively correlated with the alpha power for both low (*β*_1_ = 0.28077, R^2^ = 0.425, p = 0.0115) and high (*β*_1_ = 0.17617, R^2^ = 0.38, p = 0.0186) load condition (**Figure 11(A)**). *α*/*β* Power ratio was also found to be a significant predictor of precision in low (*β*_1_ = 0.11906, R^2^ = 0.69, p = 0.0002) and high load (*β*_1_ = 0.063459, R^2^ = 0.4, p = 0.008) conditions across lifespan (**Figure 11(B)**).

**Figure 11.**
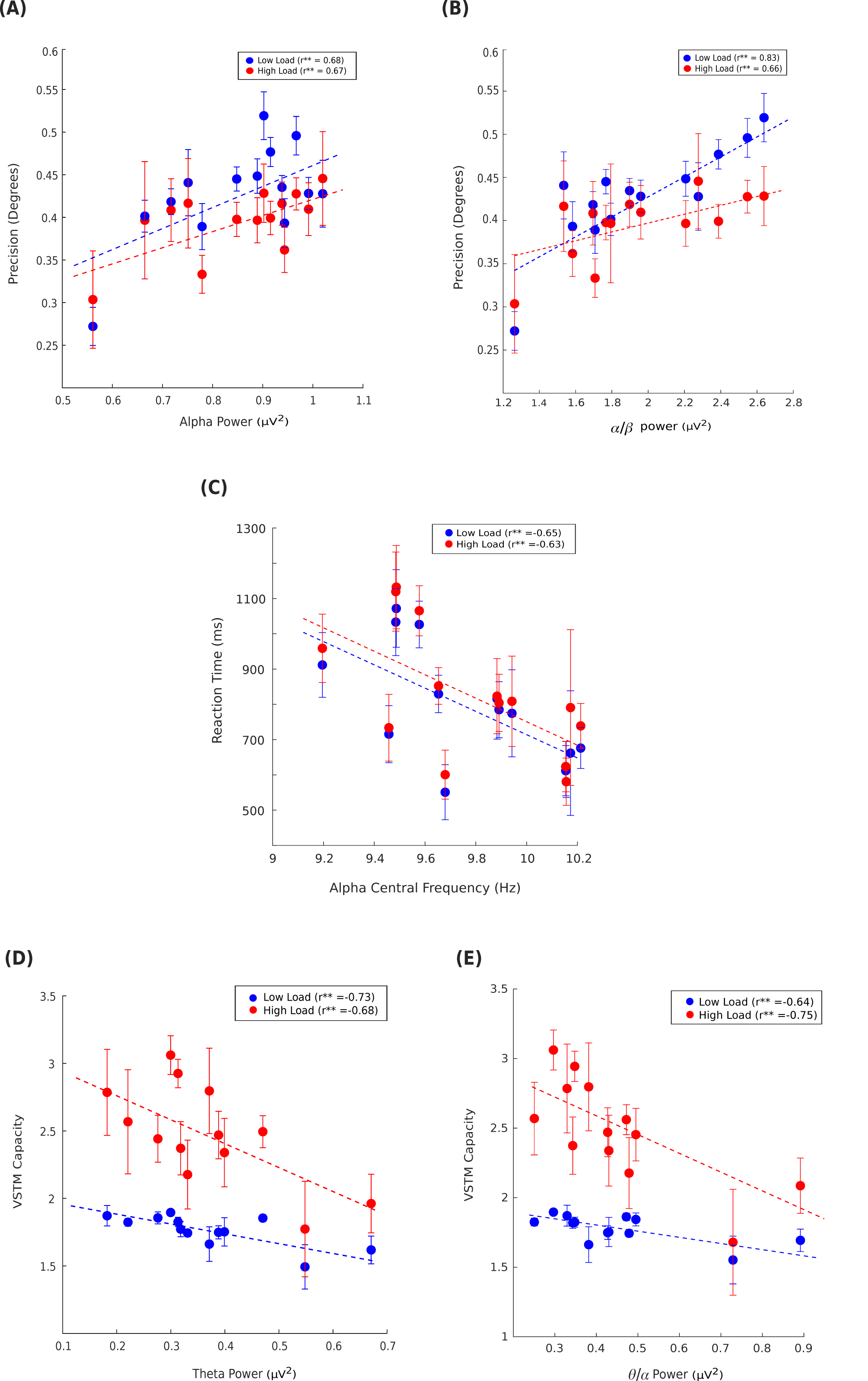
VSTM measures predicted by different oscillatory features and 1/f offset. **(A) & (B)** Precision predicted by global Alpha power and *α*/*β* Power Ratio. **(C)** Speed of Processing (RT) well predicted by global alpha speed (PAF). **(D) & (E)** VSTM capacity predicted by global theta power and *θ*/*α* Power Ratio. Age is regressed out. Low load and high load indicate set size 2 and 4 respectively. The dashed line represents the regression line. Errorbar represents SEM. ‘r’ corresponds to Pearson’s coefficient.

### 7. Speed of Processing predicted by Alpha speed

Speed of Alpha is often related to the speed of processing which is generally measured as reaction time. As we observed that speed of alpha decreases and RT increases with age, we wanted to investigate if this decrease in alpha speed affected the speed of processing in older adults. Alpha Speed significantly predicted the speed of processing for both low (*β*_1_ = -340.82, R^2^ = 0.43, p = 0.0108) and high (*β*_1_= -352.41, R^2^ = 0.39, p = 0.0158) load conditions (**Figure 11(C)**).

### 8. VSTM Capacity predicted by Theta power and ***θ***/***α*** power ratio

We found a significant negative correlation of VSTM capacity with theta power (Low Load: r = -0.729, p = 0.004, High Load: r = -0.679, p = 0.01) and *θ*/*α* power ratio (Low Load: r = -0.64, p = 0.01, High Load: r = -0.75, p = 0.001), suggesting that these two play an important role in storing items in working memory. Regression analysis also revealed a significant role of Theta power and *θ*/*α* power ratio in predicting VSTM capacity in low (Theta Power: *β*_1_ = -0.63163, R^2^ = 0.53, p = 0.0046, *θ*/*α*: *β*_1_= -1.5827, R^2^ = 0.569, p = 0.002) and High load conditions (Theta Power: *β*_1_ = -1.8862, R^2^ = 0.46, p = 0.011, *θ*/*α*: *β*_1_ = -0.35435, R^2^ = 0.42, p = 0.01) (**Figure 11(D & E)**)

## DISCUSSION

We investigate the resting-state temporal dynamics (oscillatory and non-oscillatory) and how these dynamical changes affect the cognitive and metacognitive aspects of visual short-term memory across the adult lifespan in a large cohort consisting of 280 participants. Most of the studies in aging literature have associated task-related oscillatory differences with the behavioural responses (Clark et al., 2004; Rondina et al., 2019; Proskovec et al., 2016; Cummins et al., 2007; Tóth et al., 2015). As the resting-state serves as a baseline/control for the task-related changes, it is important to characterise the dynamics in the baseline state. Consistent with the findings of Mitchell et al., 2018, we found that aging is associated with decreased Precision, memory capacity, subjective uncertainty (only in low condition) and increased reaction time in both low and high load conditions of VSTM task.

Power spectral density of the electrical activity follows the power-law distribution (1/f) which represents broadband scale-free activity of the brain (He, 2014; He et al., 2010). Most of the studies have used narrowband power analysis that presumes that spectral power implies oscillatory power, without precisely separating the 1/f aperiodic activity which in itself is dynamic and it impacts the oscillatory power which can lead to misinterpretation of the results. We approached this problem by applying a parameterization model (Voytek et al., 2020) which considers power spectrum as the combination of oscillatory peaks characterised by their Central Frequency, Power, Bandwidth measures, and aperiodic 1/f noise characterised by slope and offset. **Extended Fig. 3-5** shows the relation between 1/f slope and dominant periodic features, indicating the interdependence of these two components and the necessity to detangle these. This 1/f slope index for the noise in the brain. Aging is associated with an increase in cortical neural noise, where studies have previously used RT as a proxy for the neural noise (e.g., Creemer & Zeef, 1987; Salthouse & Lichty, 1985; Welford, 1981). Specifically, Voytek and colleagues had suggested that flattening of PSD slopes might be a hallmark of age-related cognitive decline. In support of the Neural noise hypothesis of aging, our results show that 1/f slope of the MEG spectral power increases with age (less negative), which is suggestive of increased unsynchronised neuronal activity (Hong and Rebec, 2012; Podvalny et al., 2015). Additionally, the increase in 1/f slope follows a monotonic non-linear relationship with age indicating that the rate of change in unsynchronised neural activity is not constant across adult lifespan. We observe some deviation from the normal trend for both 1/f slope and offset in age-group 60-80, which might be due to the observed increased variance in the older group. This increase in 1/f slope (increase in the local excitation/inhibition ratio), accounts for the reduction in signal to noise ratio, which in turn disrupts the fidelity of communication between the neurons and therefore it has been shown to be predictive of performance in working memory tasks (Voytek et al., 2015), N400 lexical prediction (Dave et al., 2017) and in grammar learning (Cross et al., 2020). Most importantly, we show that the RS cortical 1/f slope is predictive of cognitive performance along with the metacognitive aspect which is indirectly measured by the subjective uncertainty in visual short term memory.

Besides, aperiodic features were found to not only vary across subjects (more for elderly) but also across different sensors indicating substantial variability and idiosyncrasy. Though 1/f slope shows spatial heterogeneity in the young group, such as being less negative in the anterior sensors compared to the posterior sensors, older participants display a more homogenous distribution of less negative 1/f slope values. This global increase in noise leads to reduction of signal-to-noise ratio which would affect the information processing. Along with consideration of previous studies where 1/f slope is found to be predictive of measures in different modalities or domains, our results seem to suggest that age-related increase in neuronal noise affects global information processing which is also reflected in poorer cognitive performance of the older participants. The broadband offset shows no significant deviation with age, but significant between-group differences were observed.

In oscillatory dynamics, we observe a significant decline in peak alpha frequency (PAF) with age as shown by previous study (Sahoo et al. 2020), however how PAF relates to different aspects of behavioural responses in VSTM was not quantified systematically. This decrease in peak alpha frequency was not found to be localised to specific sensors, rather a global significant decrease was observed (see **Fig. 5(B)**). The speed of alpha is often associated with the speed of information processing therefore, higher alpha speed is needed for optimal performance in cognitive tasks (Surwillo et al., 1961) and determine the temporal resolution of visual perception (Samaha et al., 2015). **Fig. 11(C)** shows that the reaction time of the participants is well predicted by global alpha speed. Hence, Higher the speed of alpha, fast is the processing speed, lesser reaction time for younger adults. The relevance of Theta CF in determining memory capacity in a task is well known in the literature. A study by Moran et al.. 2010 shows that both slow and fast theta frequencies correlated to the high memory capacity, distributed across different networks. In the context of aging, we observe that theta CF slightly increases for older subjects as compared to younger adults which may itself affect the storing capacity. There are studies which have observed an increase in RS theta power in older adults (Klimesch et al., 1999 for review; Klass et al., 1995) others have reported theta power decrease in resting as well as in the task with age (Vlahou et al., 2014, Babiloni et al., 2006, Cummins et al., 2007, Leirer et al., 2011). However, we found an increase in theta power with age, which significantly predicted the VSTM capacity (**Fig. 6(A), 11(E)**) along with *θ*/*α* power ratio (TAR) (**Fig.8**) which also significantly increases with age. Few studies including the study by Trammell et al., 2017 which found decreased performance in RM correlated with increased TAR in old adults. Though we found substantial variability in the presence of theta power in participants. For instance, in young groups, the theta was not observed over frontal and left temporal sensors, whereas in older participants the theta power was observed only over temporal sensors.

One of the theories to explain decreased capacity is from the perspective of the orientation of Attention, the inability to ignore irrelevant items in older subjects. Recent literature suggests that Alpha plays an active inhibitory role and inhibiting the task-irrelevant information is reflected by increased alpha power. As aging is characterized by attentional difficulties, in particular, a reduced capability to inhibit irrelevant information. We found that alpha power decreases with age, particularly over occipito-parietal sensors (**Fig. 6**) which significantly predicts the precision (**Fig. 11(A)**). It plays a crucial role in suppressing irrelevant information, therefore, not being able to ignore distractions might be one of the reasons for low VSTM capacity in older adults. Though study by Vaden et al., 2012 demonstrated that older people do not use alpha power suppression to inhibit distractor’s information.

We observed an increase in Beta power with age which is well reported in the literature, generally associated with the movement-related activity (Ishii et al.,2017; Sahoo et al., 2020) but we also observed a significant decrease in beta peak frequency with age (generally found in depression and other psychological disorders’ patients in open-eye condition; Roohi-Azizi et al., 2017). Particularly, the change was more localised to the central-parietal sensors. Only Beta bandwidth was found to increase with age which indicates that variability of beta frequency becomes more or it spreads out more. This increase was mostly observed in the central and temporal sensors.

A study by Griffiths et al., 2019 suggested that task-induced decrease in *α*/*β* power is a proxy for reductions in noise correlations and rather, decreased *α*/*β* power provides a favourable (i.e., reduced noise) condition in which another mechanism can allow the signal representation in a task. Though the time-scale in our study is different compared to them, our result shows that a decrease in *α*/*β* power is a significant predictor of precision (**Fig. 11 (B)**). After regressing out the age factor, the increase in 1/f slope is negatively correlated with the *α*/*β* power which suggests that *α*/*β* cannot serve as a proxy for noise (**Extended fig. 9-4**). This increase in 1/f slope might be the reason for the decrease in *α*/*β* power, which in turn decreases the precision.

Our results suggest that the age-associated global change in noisy baseline affects the global information processing and link to cognitive decline in the performance of old elderly in a short-term working memory task. Specifically, an increase in slope with age affects the speed of information processing, cognitive capacity, precision and metacognitive awareness (all behavioural measures used in this study). In contrast, oscillatory features of different frequency bands which are crucially impacted by the baseline shift in the global noisy background relate to more local processing and selective behavioural measures in VSTM task. On that account, global increase in noise (indexed by 1/f slope) seems to impact distributed processes of cognition whereas oscillatory features mediate localised processing, involved in a particular cognitive task (**Figure 12**).

**Figure 12.**
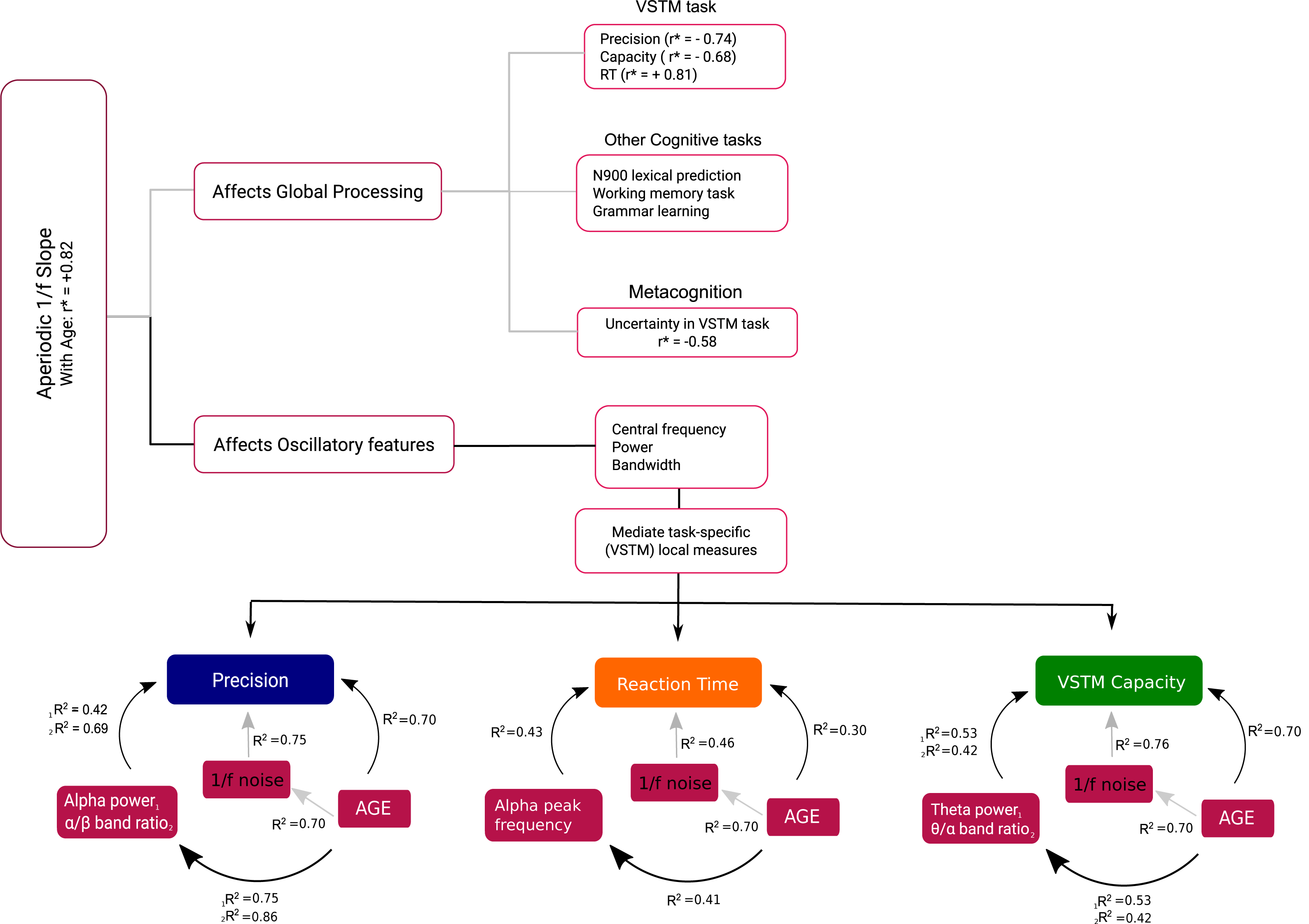
Aperiodic 1/f activity affects global processingand oscillatory features. Aperiodic 1/f slope increases globally which can be observed in performance in different cognitive tasks (affects global processing). It also significantly affects oscillatory features which are more task-specific.

Conclusively, we suggest that the differences we observe in the task-related functional changes (across age groups) are relative to their baseline. RS dynamics, for instance the power of oscillation, with any incoming visual stimuli will change relative from its power baseline and with age, these baseline gets shifted, which are reflected in the cognitive performance. Hence, the differences that we observe in the ‘task-related functional changes’ across different age groups might be due to the change in the relative spectral features’ baseline with age.

An important limitation of our study is that we have only tested our hypothesis in VSTM task, therefore, further investigation is needed relating the RS 1/f noise with the performance in different cognitive tasks and also how these patterns change from their respective baseline when the task is being performed. Another limitation was posed by the Cam-CAN dataset, because of the presence of harmonics of lower frequencies in higher frequencies, we were not able to systematically analyse the effect of 1/f noise on gamma-band. Lastly, we still don’t know which sources are responsible for this 1/f baseline shift and can be considered for future work, where source reconstruction and applying the model on the source level spectrum might give more clarity about the sources. Despite these limitations, our study characterises the periodic and aperiodic temporal dynamics of the sensor-level neural activity across the adult lifespan with a large cohort and reasoned how the shifted 1/f noise baseline affects cognition.

## Data/code availability statement

The datasets generated during and/or analysis pipelines used during the current study are available from the corresponding author on reasonable request.

## Acknowledgements

This study was supported by NBRC Core funds, Ramalingaswami Fellowships (Department of Biotechnology, Government of India) to DR (BT/RLF/Re-entry/07/2014) and AB (BT/RLF/Re-entry/31/2011) and Innovative Young Biotechnologist Award (IYBA) to AB (BT/07/IYBA/2013). AB also acknowledges the support of the Centre of Excellence in Epilepsy and MEG. DR was also supported by SR/CSRI/21/2016 extramural grant from the Department of Science and Technology (DST) Ministry of Science and Technology, Government of India. Data collection and sharing for this project was provided by the Cambridge Centre for Aging and Neuroscience (CamCAN). CamCAN funding was provided by the UK Biotechnology and Biological Sciences Research Council (grant number BB/H008217/1), together with support from the UK Medical Research Council and University of Cambridge, UK. In accordance with the data usage agreement for CAMCAN dataset, the article has been submitted as open access.

---

**Figure 3-1.**
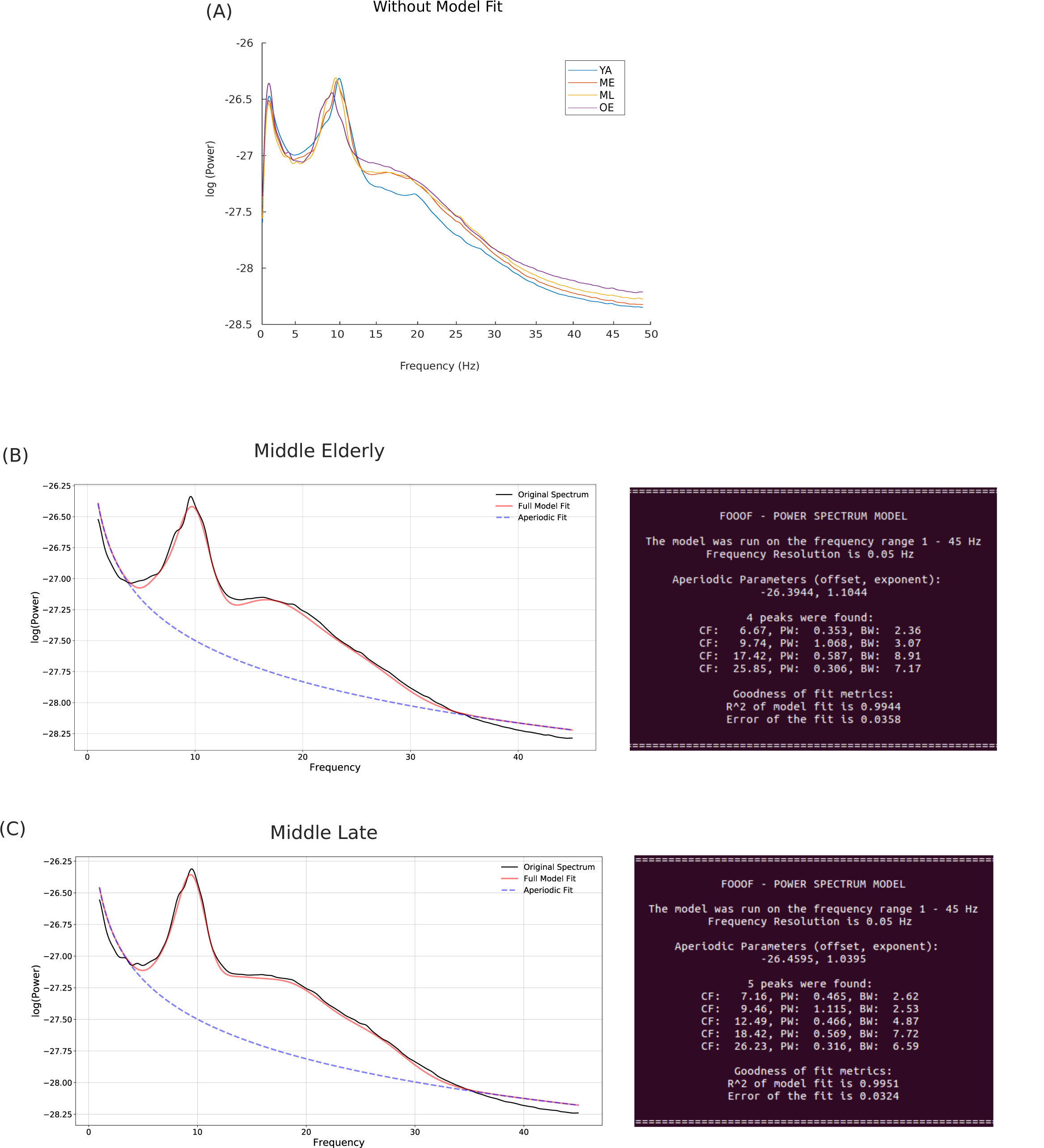
(A) Power spectrum without model fitting (B) & (C) FOOOF-Power spectrum model for ME and ML age groups, along with its statistics

**Figure 3-2.**
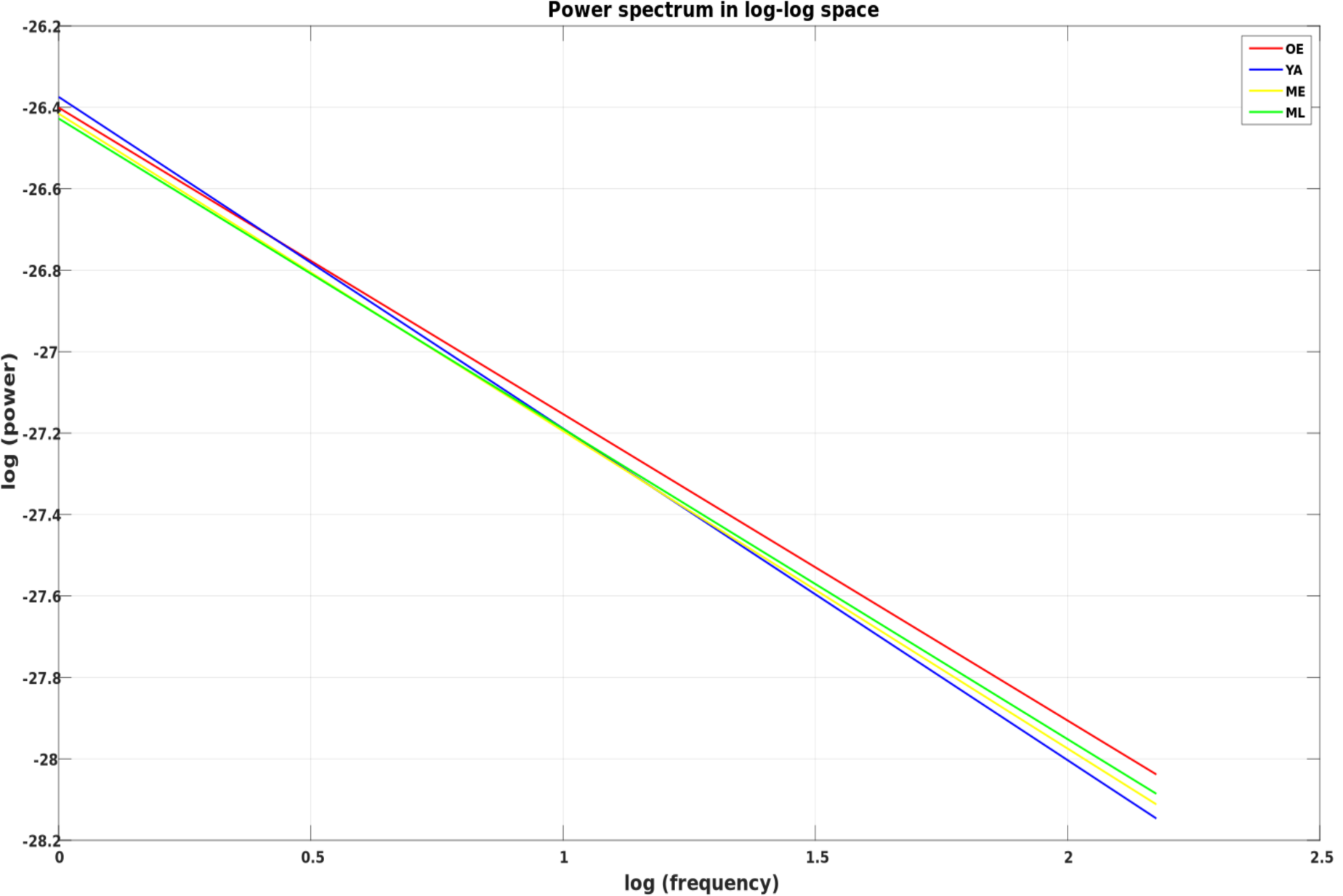
Power Spectrum in log-log space

**Figure 3-3.**
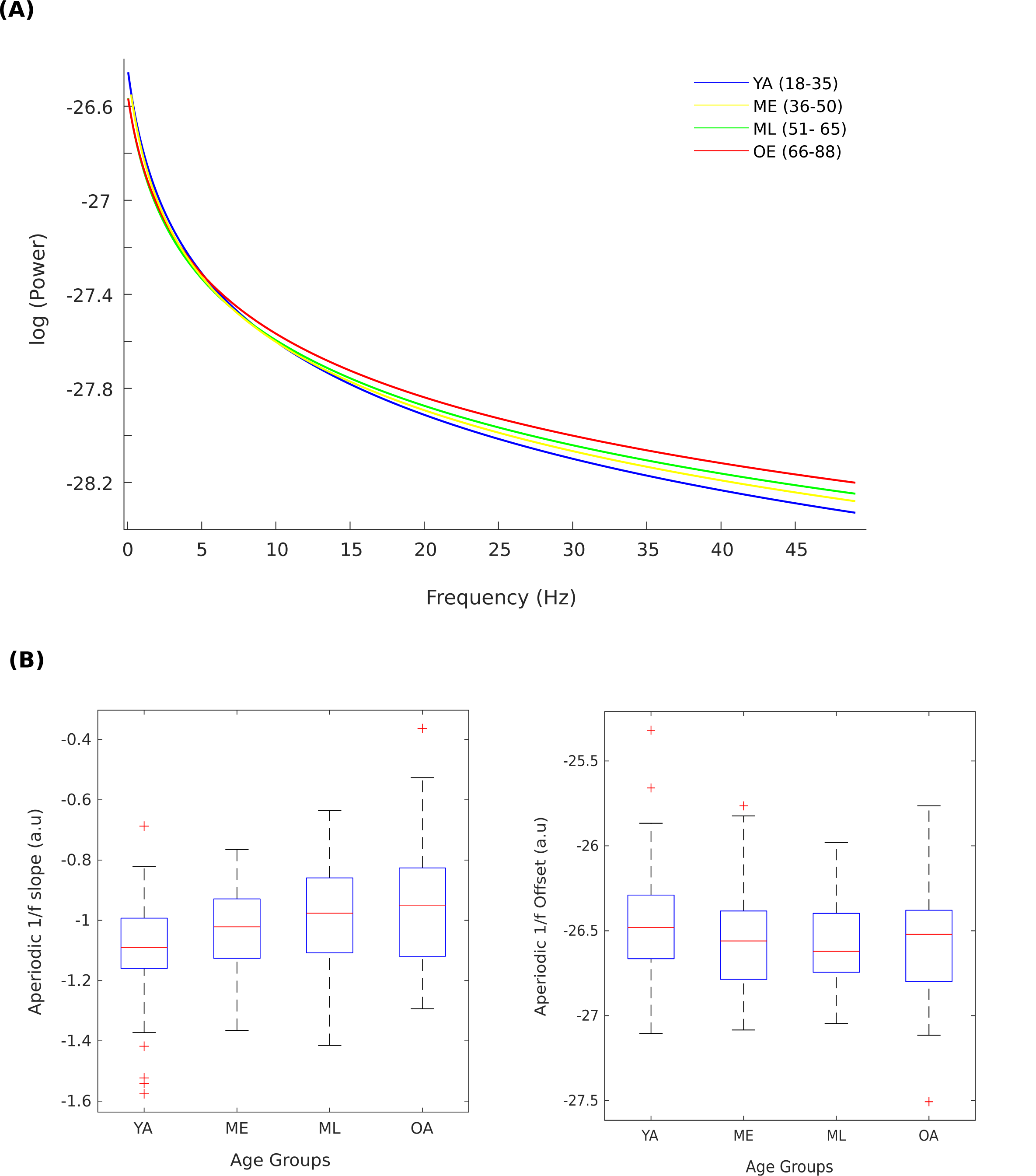
(A) Aperiodic component of different age groups (B) Boxplot of aperiodic features aross different age groups derived from the component.

**Figure 3-4.**
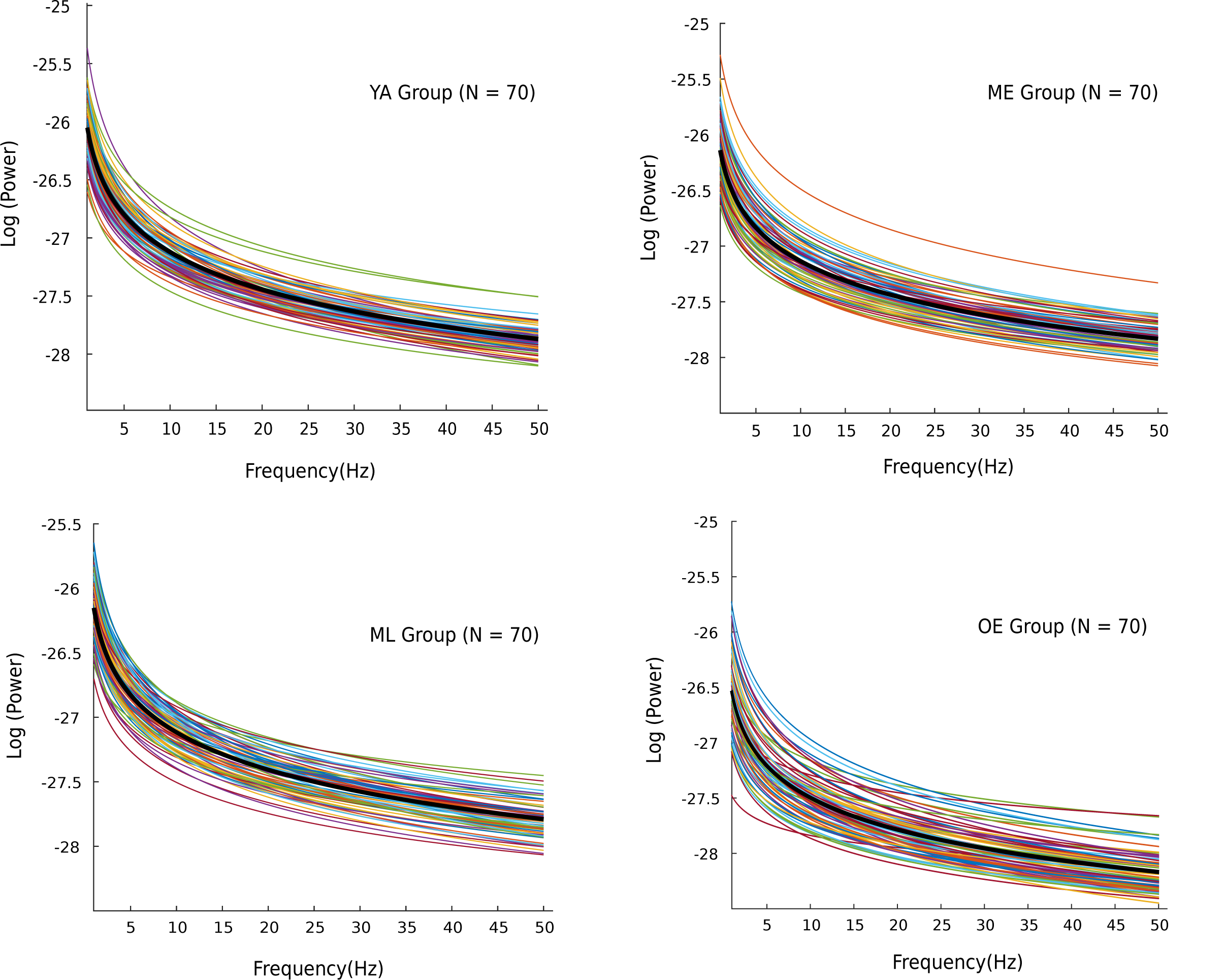
Aperiodic component vary across subjects in each age group

**Figure 3-5.**
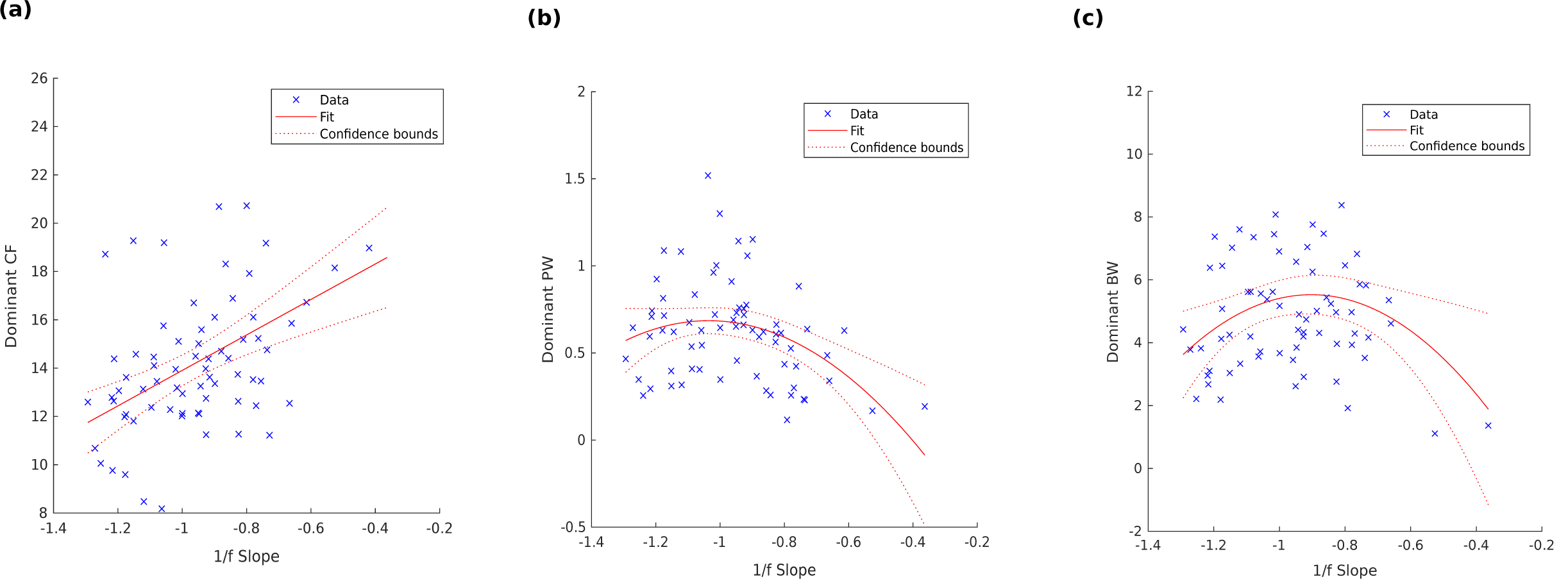
Regression plot for 1/f aperiodic slope with periodic components of the dominant oscillations.

**Figure 5-1.**
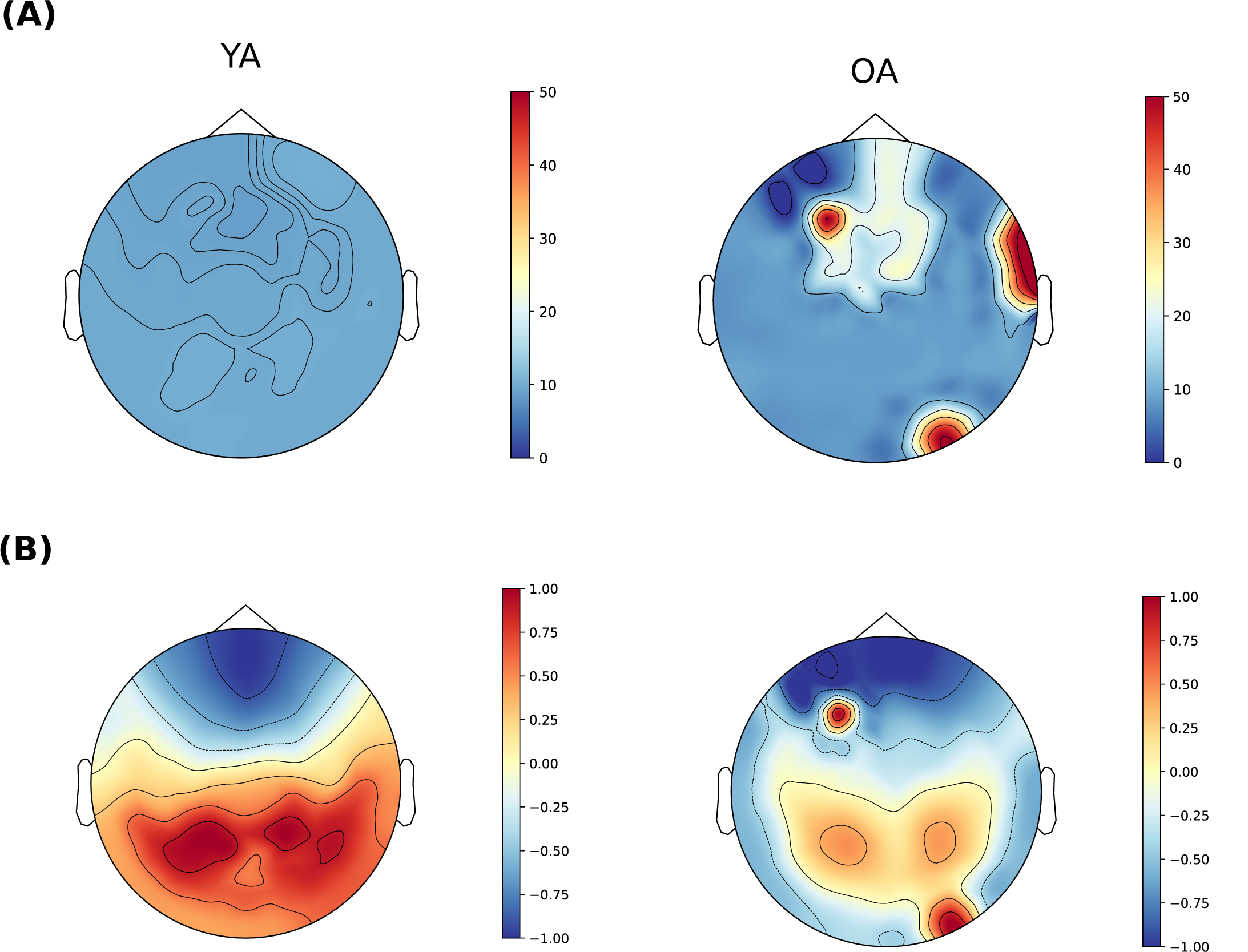
Dominant frequency (A) and respective power (B) for YA and OA

**Figure 6-1.**
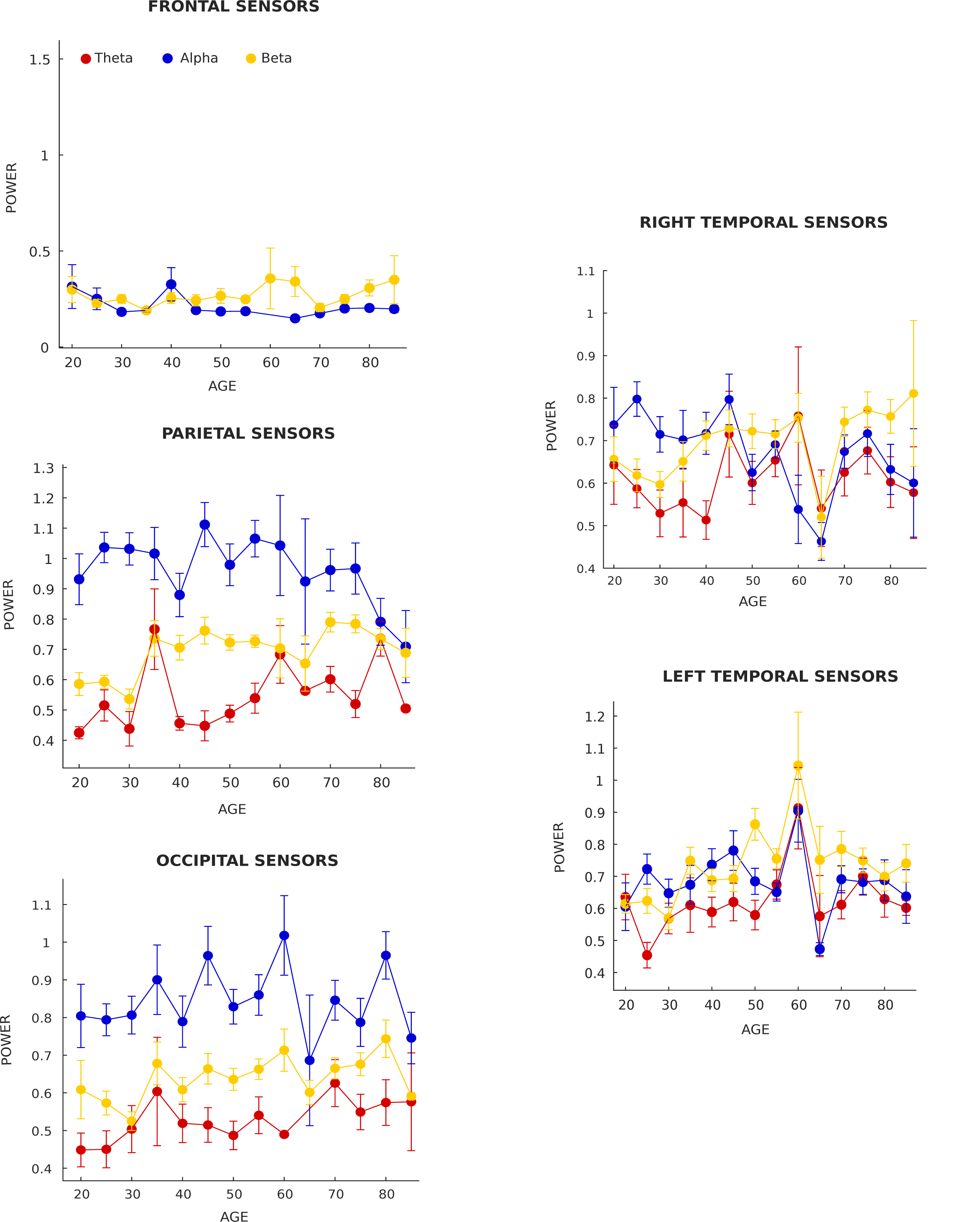
Frequency specific power as a functionof age across different sensors. For Frontal sensors, theta power is not shown as only few age groups showed peaks in theta range. Error bar represents SEM

**Figure 7-1.**
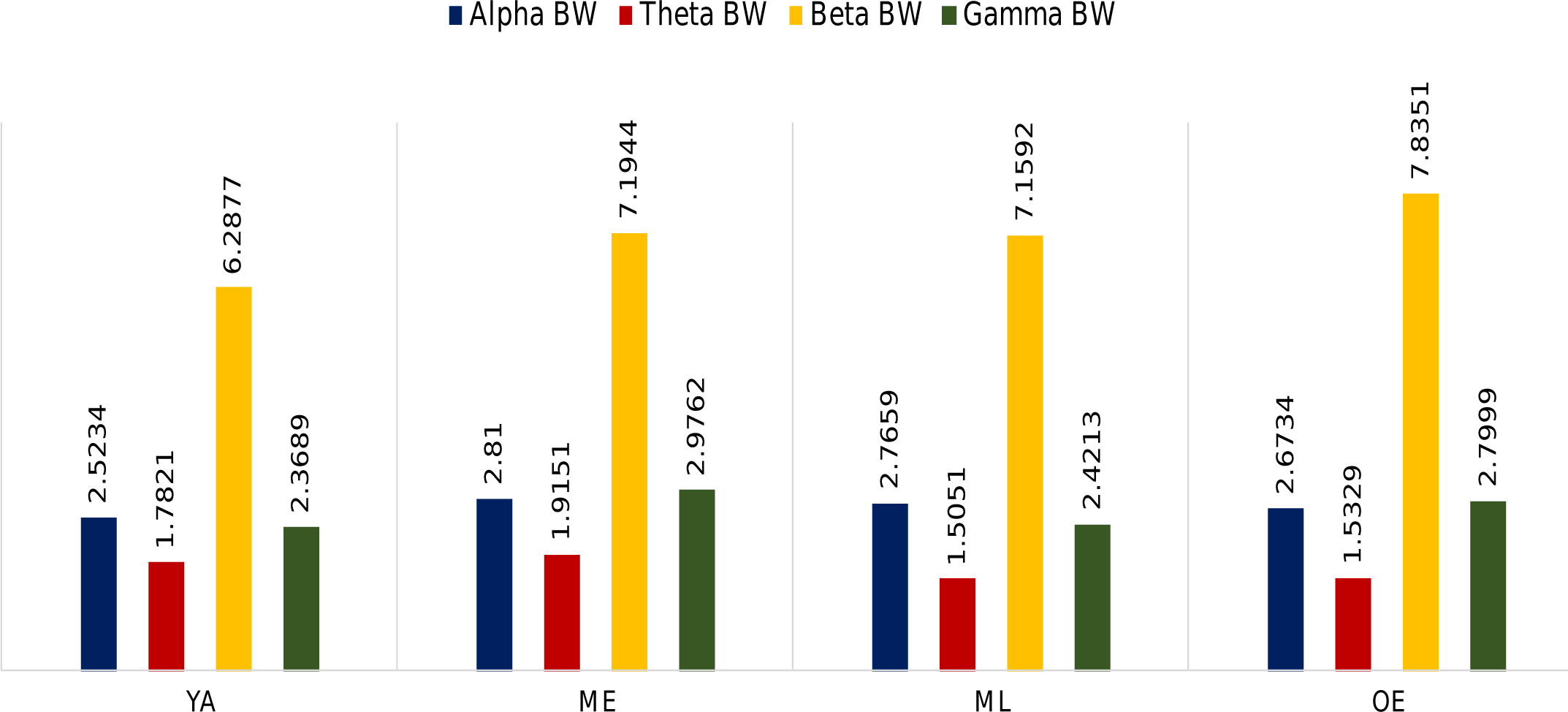
Frequency specific bandwidth of different age groups

**Figure 7-2.**
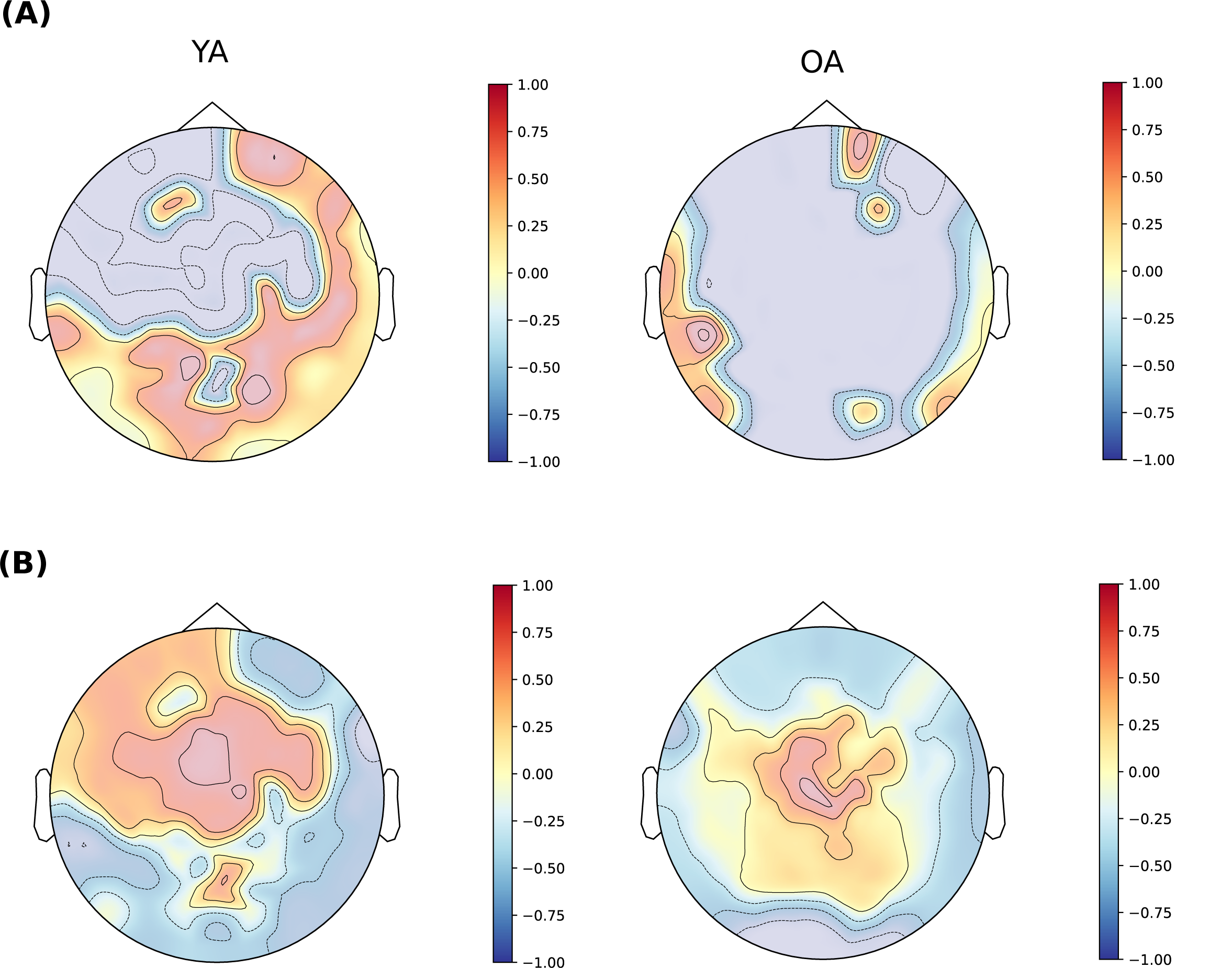
Spatial topography of Theta Bandwidth(A) and Alpha Bandwidth (B) for YA and OA

**Figure 8-1.**
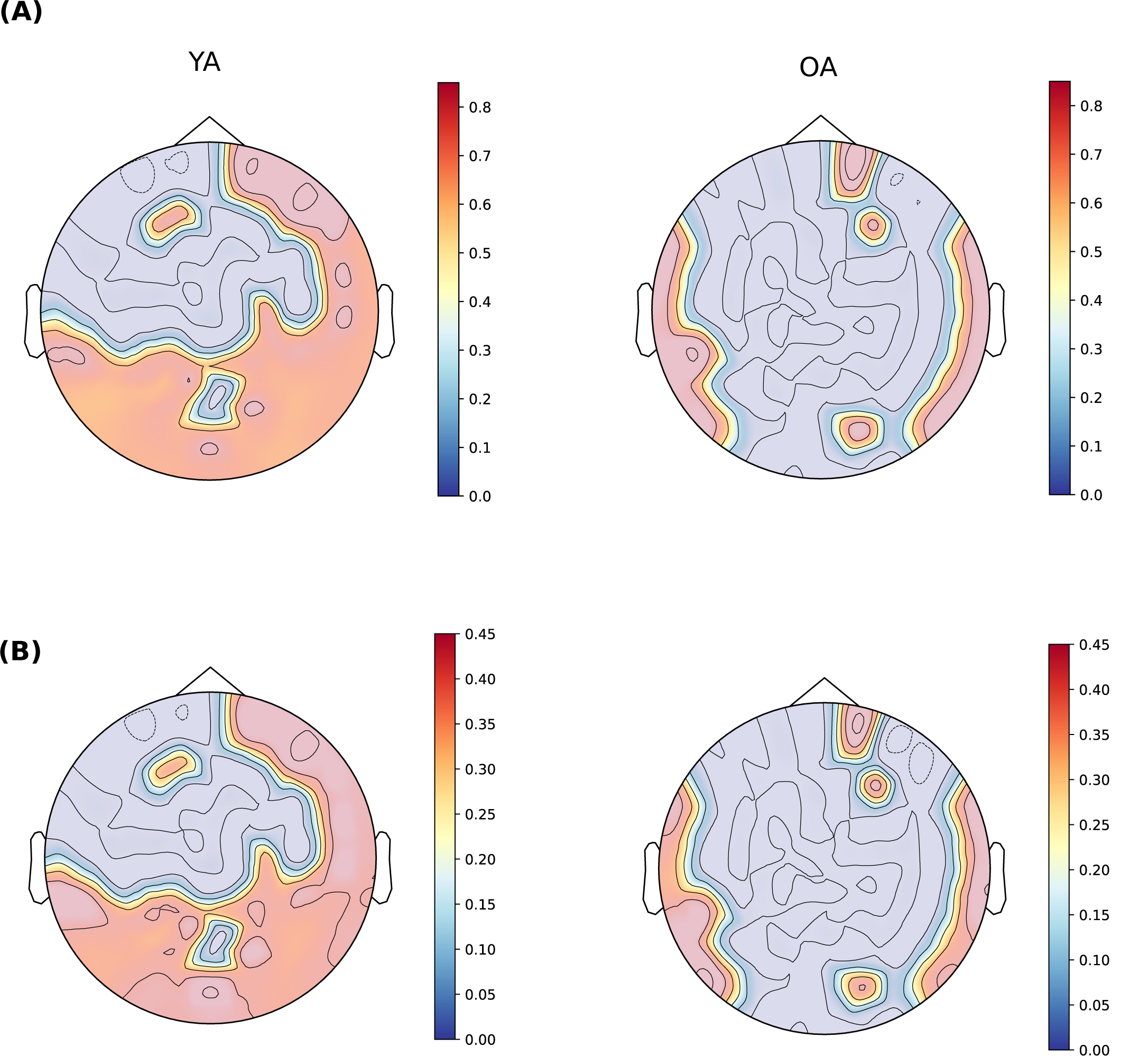
**(A)** Theta/Alpha peak frequency ratio and **(B)** Theta/Beta peak freqeuncy ratio for YA and OA

**Figure 8-2.**
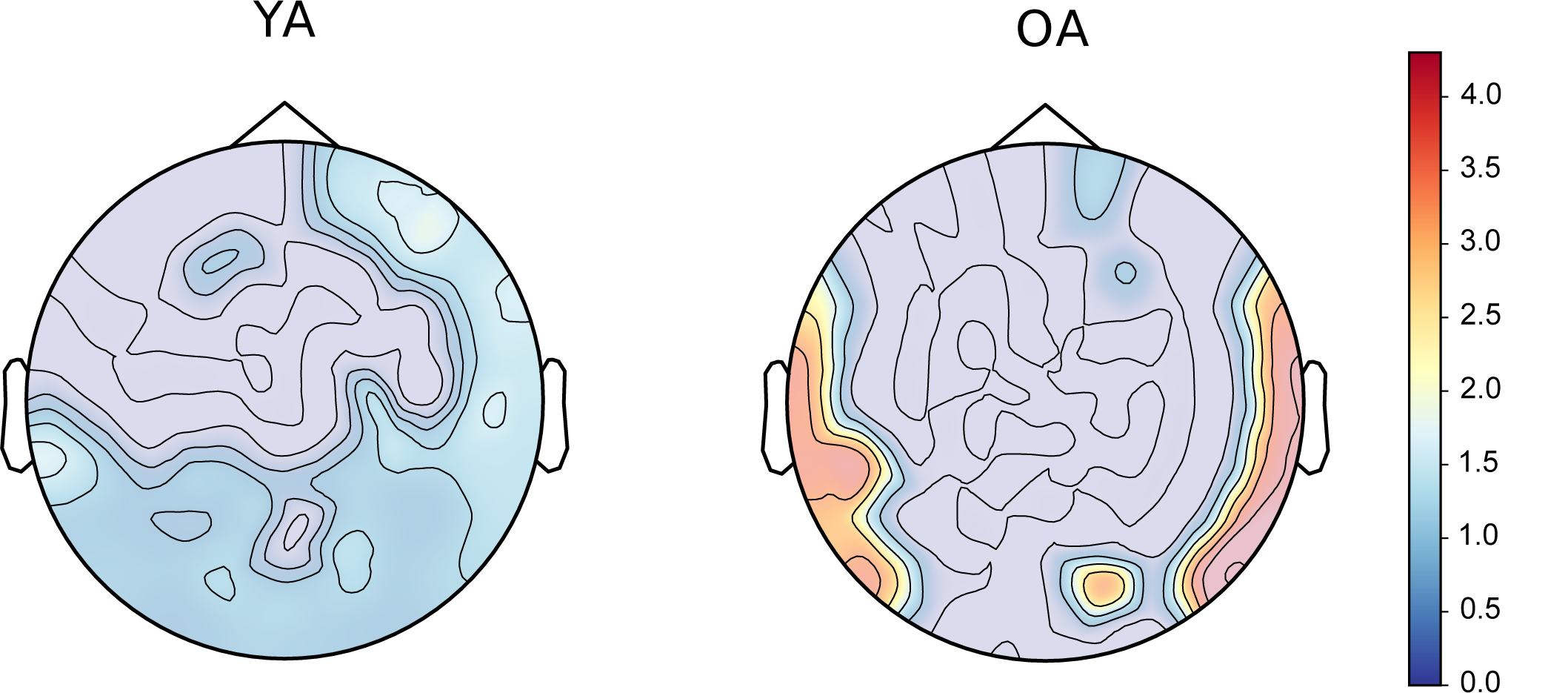
Theta/Beta PW ratio as a function of age

**Figure 8-3.**
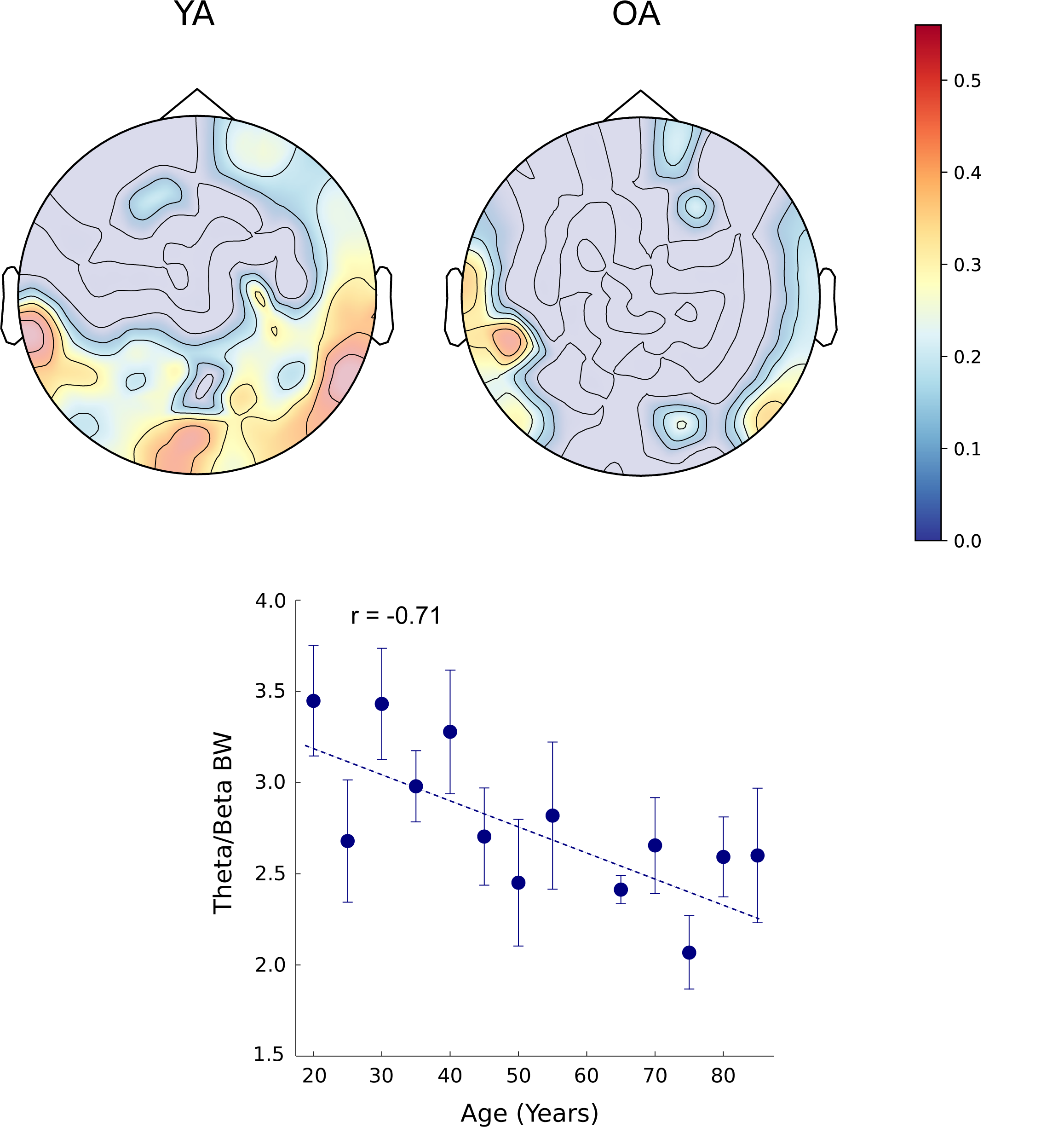
Theta/Beta BW as a function of age

**Figure 8-4.**
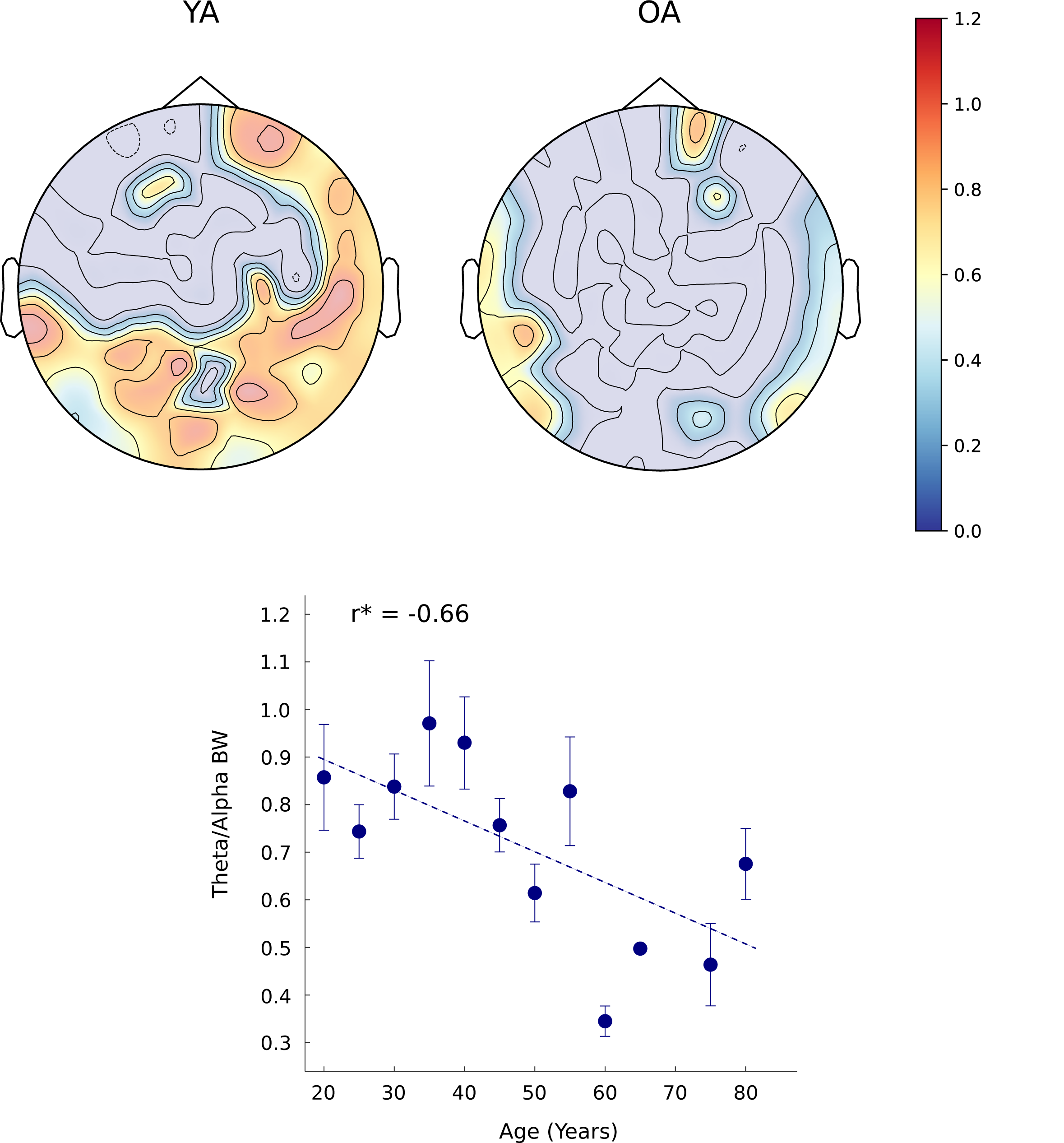
Theta/Alpha BW as a function of age

**Figure 8-5.**
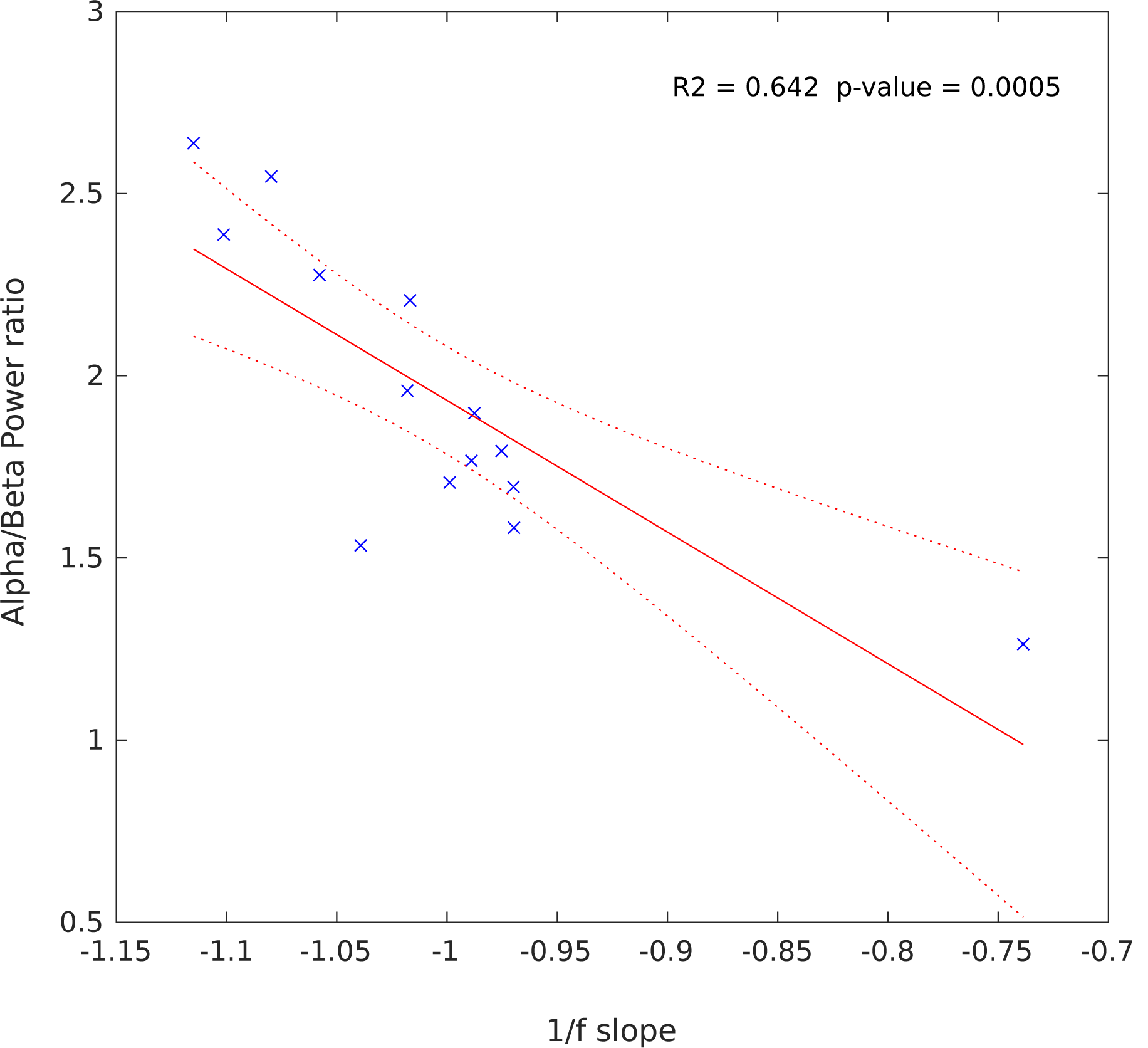
Regression model for 1/f slope with Alpha/Beta power ratio

**Figure 9-1.**
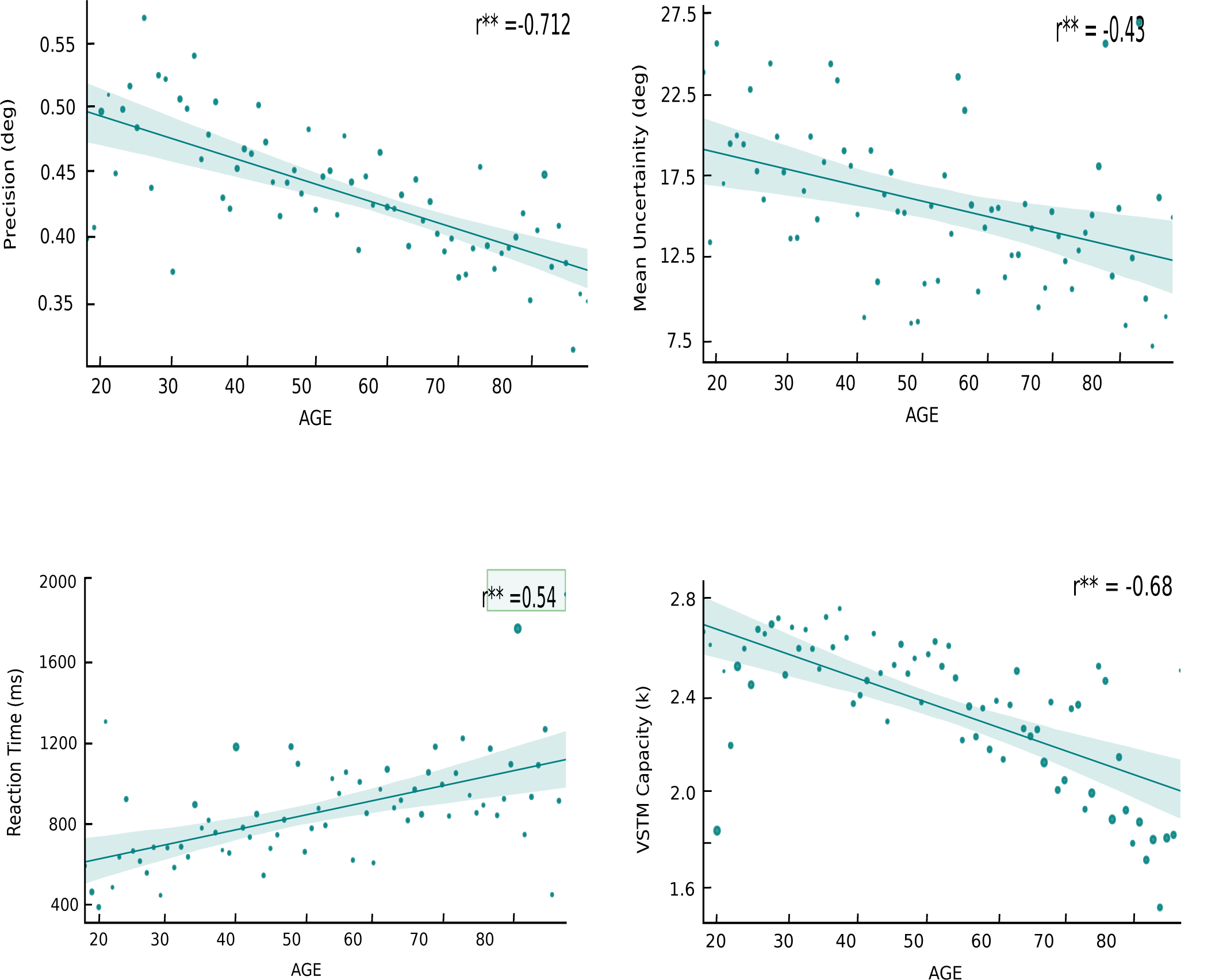
VSTM measures as a function of age. Particiapnts with same age were grouped togther (dots), size of the dot represents the SEM of the group. Shaded area is the 95% confidence interval.

**Figure 9-2.**
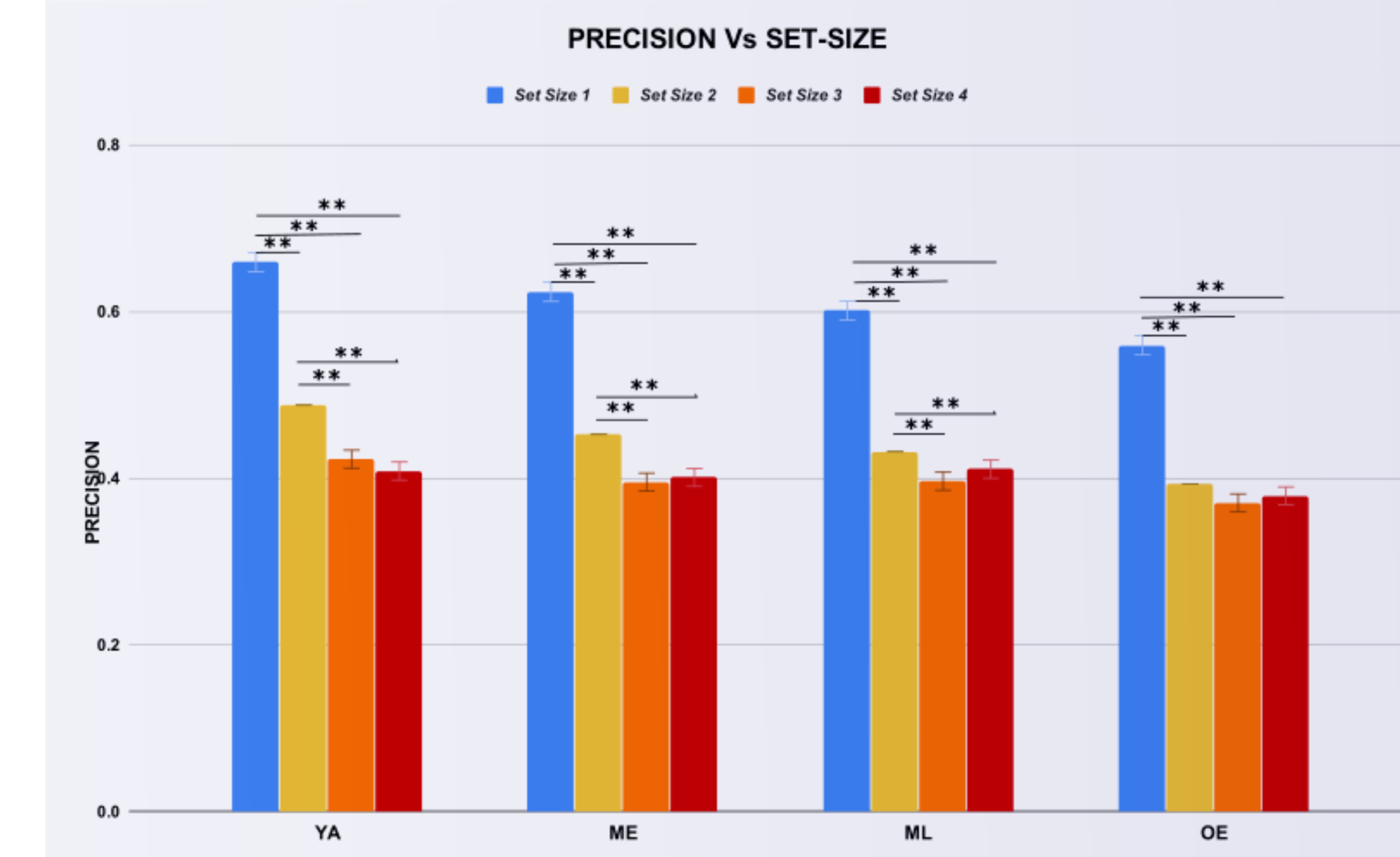
Precision across age groups and set size

**Figure 9-3.**
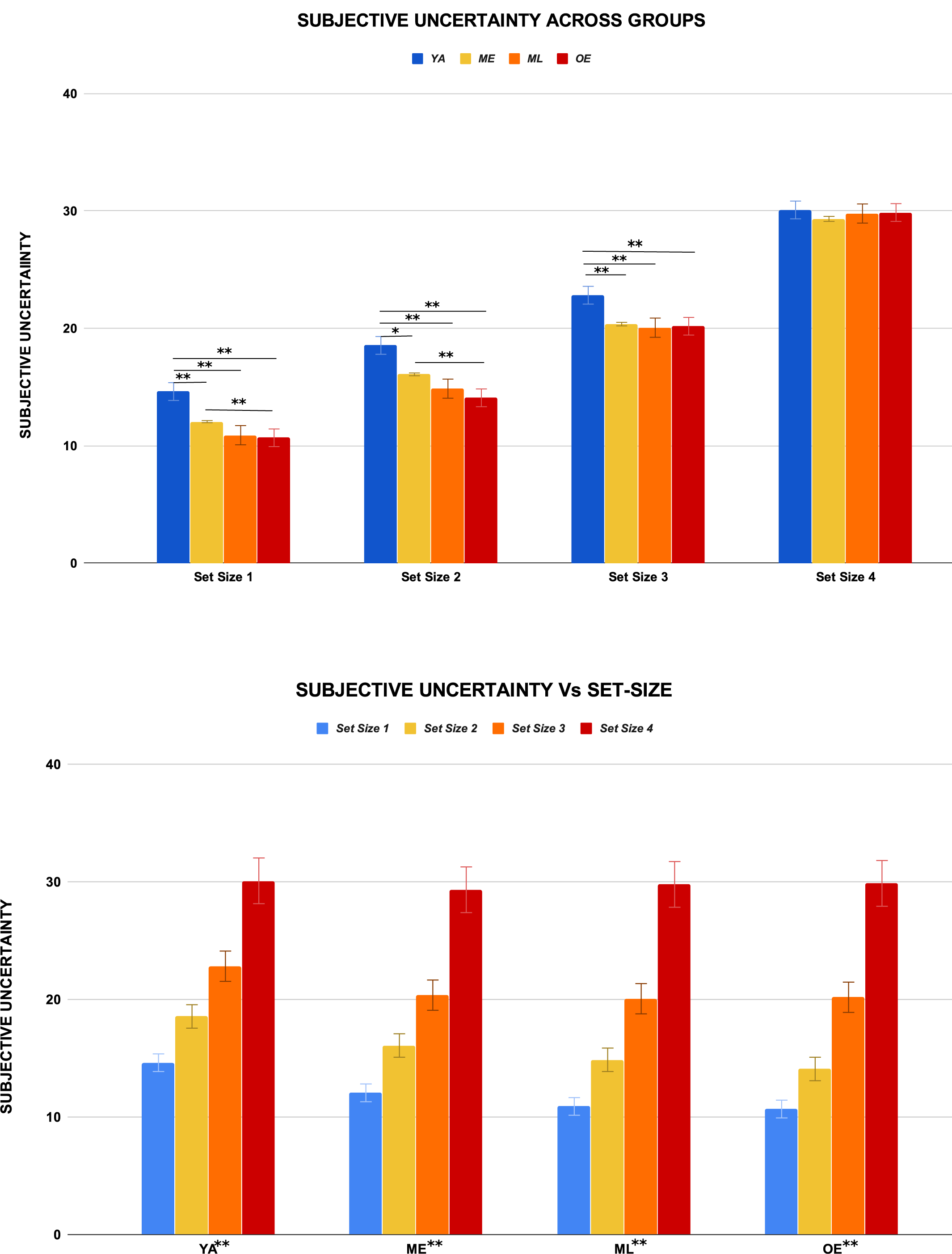
Uncertainty across groups and set size

**Table 8-1:** Regression table for global frequency band-ratios with Age. F-value, Beta coefficient, Goodness of fit and significance of the model is reported.

| Explanatory Variable | Response Variable | F-value | Beta1 | p-value | R<sup>2</sup> |
| --- | --- | --- | --- | --- | --- |
| Age | $\alpha/\beta$ PW | 77 | -0.018116 | 1.44e<sup>-06</sup> | 0.86 |
| | $\alpha/\beta$ CF | 8.7 | -0.005913 | 0.0054 | 0.62 |
| | $\theta/\beta$ BW | 5.31 | -0.001542 | 0.03 | 0.3 |
| | $\theta/\alpha$ PW | 7.2 | +0.00576 | 0.02 | 0.39 |

**Table 9-1:**
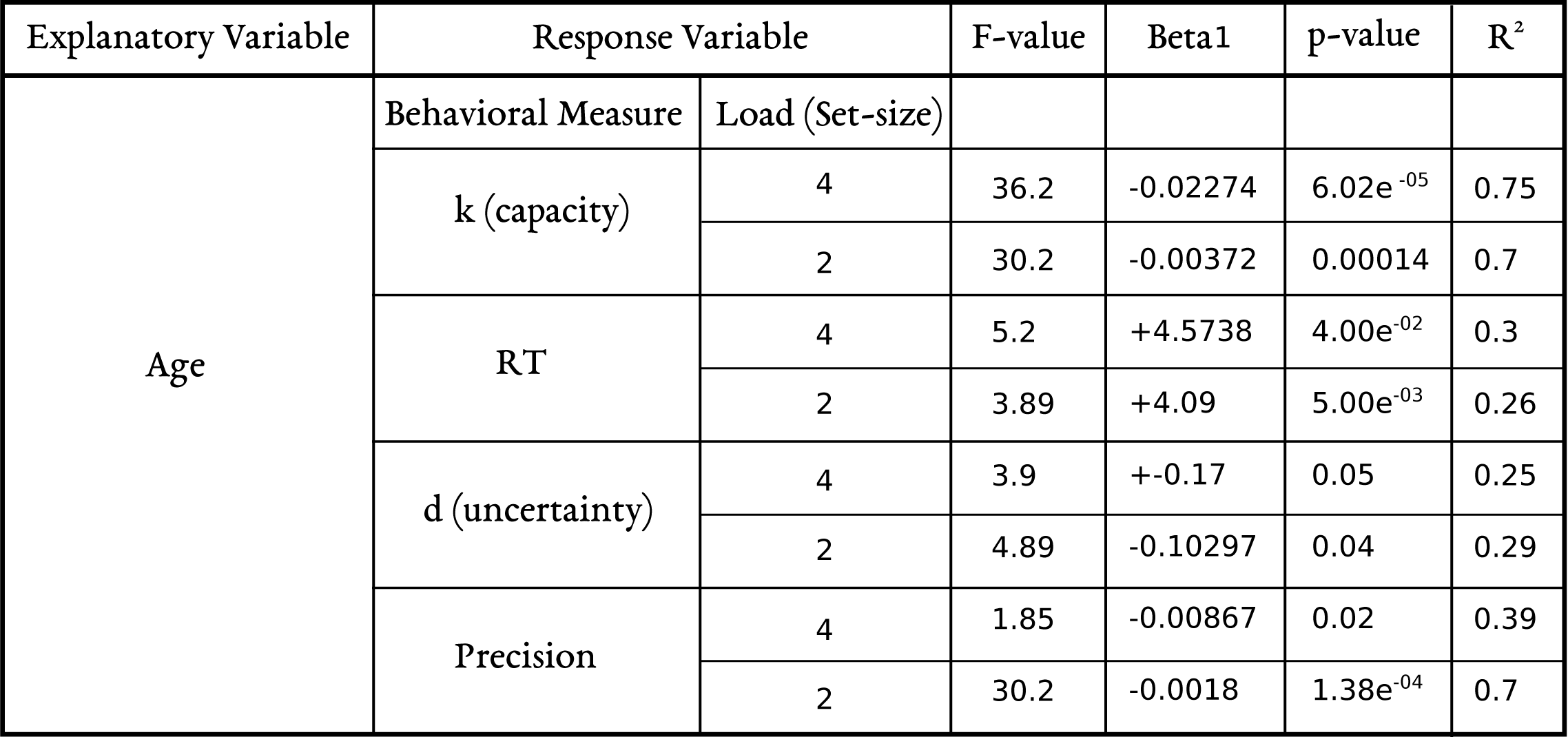
Regression table for VSTM measures with Age. F-value, Beta coeficient, Goodness of fit and significance of the model is reported.

**Table 10-1:** Regression table for VSTM measures with aperiodic slope. F-value, Beta coeficient, Goodness of fit and significance of the model is reported.

| Explanatory Variable | Response Variable |  | F-value | Beta1 | p-value | R<sup>2</sup> |
| --- | --- | --- | --- | --- | --- | --- |
| Aperiodic 1/f Slope | Behavioral Measure | Load (Set-size) |  |  |  |  |
|  | k (capacity) | 4 | 25.9 | -6.2486 | 0.0003 | 0.702 |
|  |  | 2 | 2.98 | -0.75038 | 0.112 | 0.2 |
|  | RT | 4 | 10.6 | +2666.3 | 0.0076 | 0.49 |
|  |  | 2 | 9.59 | +2336.3 | 0.0102 | 0.466 |
|  | d (uncertainty) | 4 | 0.25 | -46.1 | 0.02 | 0.35 |
|  |  | 2 | 5.5 | -48.817 | 0.0388 | 0.33 |
|  | Precision | 4 | 1.52 | -0.10677 | 0.24 | 0.11 |
|  |  | 2 | 36.9 | -0.44753 | 5.52e<sup>-05</sup> | 0.75 |

**Table 10-2:** Regression table for VSTM measures with Aperiodic 1/f ofiset. F-value, Beta coeficient, Goodness of fit and significance of the model is reported.

| Explanatory Variable | Response Variable |  | F-value | Beta1 | p-value | R<sup>2</sup> |
| --- | --- | --- | --- | --- | --- | --- |
| Aperiodic 1/f<br>Offset | Behavioral Measure | Load (Set-size) |  |  |  |  |
|  | k (capacity) | 4 | 2.06 | +2.3507 | 0.179 | 0.158 |
|  |  | 2 | 1.53 | +0.4515 | 0.241 | 0.122 |
|  | RT | 4 | 8.26 | -1979.2 | 0.01 | 0.429 |
|  |  | 2 | 10.1 | -1880.9 | 0.008 | 0.47 |
|  | d (uncertainty) | 4 | 0.168 | -14.374 | 0.69 | 0.015 |
|  |  | 2 | 0.559 | -14.762 | 0.47 | 0.04 |
|  | Precision | 4 | 5.2e<sup>-05</sup> | -0.0009 | 0.99 | 4.7e<sup>-06</sup> |
|  |  | 2 | 0.658 | -0.1406 | 0.52 | 0.05 |

**Table 11-1:** Regression table for specific oscillatory features with VSTM measures. F-value, Beta coeficient, Goodness of fit and significance of the model is reported.

| Explanatory Variable | Response Variable |  | F-value | Beta1 | p-value | R<sup>2</sup> |
| --- | --- | --- | --- | --- | --- | --- |
| Alpha CF | Behavioral Measure | Load (Set-size) |  |  |  |  |
|  | RT | 4 | 7.89 | -352.41 | 0.0158 | 0.39 |
|  |  | 2 | 9.08 | -340.82 | 0.0108 | 0.43 |
| Alpha PW | Precision | 4 | 7.4 | +0.1761 | 0.0186 | 0.38 |
|  |  | 2 | 8.89 | +0.2807 | 0.0115 | 0.42 |
| Theta PW | k(capacity) | 4 | 9.46 | -1.8862 | 0.0106 | 0.46 |
|  |  | 2 | 12.5 | -0.63163 | 0.0046 | 0.53 |

**Table 11-2:** Regression table for **α*/*β** CF with VSTM measures. F-value, Beta coeficient, Goodness of fit and significance of the model is reported.

| Explanatory Variable | Response Variable |  | F-value | Beta1 | p-value | R<sup>2</sup> |
| --- | --- | --- | --- | --- | --- | --- |
| $\alpha/\beta$ CF | Behavioral Measure | Load (Set-size) | | | | |
|  | k (capacity) | 4 | 3.09 | -6.3816 | 0.1 | 0.2 |
|  |  | 2 | 0.63 | +0.6699 | 0.44 | 0.05 |
|  | RT | 4 | 0.8 | -1510.3 | 0.38 | 0.06 |
|  |  | 2 | 1.96 | -2071.3 | 0.18 | 0.14 |
|  | d (uncertainty) | 4 | 0.58 | -53.371 | 0.46 | 0.04 |
|  |  | 2 | 0.2 | +20.287 | 0.61 | 0.02 |
|  | Precision | 4 | 6.57 | -0.7578 | 0.02 | 0.35 |
|  |  | 2 | 6.33 | +25.304 | 0.01 | 0.53 |

**Table 11-3:** Regression table for **α*/*β** PW with VSTM measures. F-value, Beta coeficient, Goodness of fit and significance of the model is reported.

| Explanatory Variable | Response Variable |  | F-value | Beta1 | p-value | R<sup>2</sup> |
| --- | --- | --- | --- | --- | --- | --- |
| $\alpha/\beta$ PW | Behavioral Measure | Load (Set-size) | | | | |
|  | k (capacity) | 4 | 15.6 | +0.7898 | 0.06 | 0.56 |
|  |  | 2 | 0.12 | -0.0219 | 0.7 | 0.01 |
|  | RT | 4 | 5.96 | -255.71 | 0.6 | 0.33 |
|  |  | 2 | 3.18 | -188.68 | 0.09 | 0.21 |
|  | d (uncertainty) | 4 | 1.98 | +6.9368 | 0.18 | 0.14 |
|  |  | 2 | 5.06 | +5.6266 | 0.055 | 0.3 |
|  | Precision | 4 | 9.69 | +0.0634 | 0.008 | 0.4 |
|  |  | 2 | 27.3 | +0.1191 | 0.0002 | 0.69 |

**Table 11-4:** Regression table for **α*/*β** PW with VSTM measures. F-value, Beta coeficient, Goodness of fit and significance of the model is reported.

| Explanatory Variable | Response Variable |  | F-value | Beta1 | p-value | R<sup>2</sup> |
| --- | --- | --- | --- | --- | --- | --- |
| *θ/α* PW | Behavioral Measure | Load (Set-size) |  |  |  |  |
|  | k (capacity) | 4 | 14.5 | -1.5827 | 0.002 | 0.57 |
|  |  | 2 | 7.96 | -0.3543 | 0.01 | 0.42 |
|  | RT | 4 | 1.25 | +371.2 | 0.288 | 0.10 |
|  |  | 2 | 2.04 | +397.1 | 0.181 | 0.16 |
|  | d (uncertainty) | 4 | 0.25 | +9.321 | 0.652 | 0.01 |
|  |  | 2 | 0.26 | -6.3127 | 0.66 | 0.01 |
|  | Precision | 4 | 0.02 | +0.013 | 0.878 | 0.002 |
|  |  | 2 | 0.15 | -0.0436 | 0.86 | 0.02 |

